# High-resolution analysis of the pneumococcal transcriptome under a wide range of infection-relevant conditions

**DOI:** 10.1101/283739

**Authors:** Rieza Aprianto, Jelle Slager, Siger Holsappel, Jan-Willem Veening

## Abstract

*Streptococcus pneumoniae* is an opportunistic human pathogen that typically colonizes the nasopharyngeal passage and causes lethal disease in other host niches, such as the lung or the meninges. The expression and regulation of pneumococcal genes at different life-cycle stages, such as commensal or pathogenic, are not entirely understood. To chart the transcriptional responses of *S. pneumoniae*, we used RNA sequencing to quantify the transcriptome under 22 different infection-relevant conditions. The transcriptomic data demonstrated a high level of dynamic expression and, strikingly, all annotated pneumococcal genomic features were expressed in at least one of the studied conditions. By computing the correlation values of every pair of genes across all studied conditions, we created a co-expression matrix that provides valuable information on both operon structure and regulatory processes. The co expression data is highly consistent with well-characterized operons and regulons, such as the PyrR, ComE and ComX regulons, and has allowed us to identify a new member of the competence regulon. Finally, we created an interactive data center named PneumoExpress (https://veeninglab.com/pneumoexpress) that enables users to access the expression data as well as the co-expression matrix in an intuitive and efficient manner, providing a valuable resource to the pneumococcal research community.

## INTRODUCTION

*Streptococcus pneumoniae* (the pneumococcus) is an opportunistic human pathogen with a high carriage rate in children, immunocompromised individuals and the elderly. The pneumococcus accounts for the majority of all mortality related to lower respiratory tract infections (LRTIs), single-handedly placing LRTIs as the deadliest communicable disease (1). Additionally, LRTIs are the second principal cause for loss of healthy life (2), with young children and the elderly especially susceptible to pneumococcal pneumonia (3, 4). In addition to lung infection, *S. pneumoniae* is responsible for other lethal infections, such as sepsis, the presence of the pneumococcus in the blood, and meningitis, the presence of pneumococcus in the cerebrospinal fluid (CSF, (5, 6)). Complicating matters, the pneumococcus is part of the typical microbiota of the respiratory tract (7–9), with four in five young children (< 5 years), and one in three adults carrying the bacterium (10, 11).

These various environments encountered by the pneumococcus demand highly adaptive and flexible regulation. Inter-host transmission, for example, involves the switch from conditions in the human nasopharynx to airborne or surface-associated droplets. Here, the bacteria must survive a lower temperature, desiccation and oxygenated air (12). In addition, sites of colonization and infections inside the host are equally challenging with varying acidities and differing levels of oxygen and carbon dioxide, diverse temperatures and a scarcity of carbon sources (13), not to mention the actions of the innate and adaptive immune system. In addition, the nasopharyngeal passage can be crowded with multiple strains of *Streptococcal* species, other bacteria, fungi and phages (13). There, the occupants are competing for resources, including location and nutrients. Thus, it is not surprising that a significant part of described pneumococcal virulence factors function as adherence factors or adhesins, which typically bind to host surface-exposed molecules like proteins and sugars to form a stable anchor onto the colonization site. Additionally, the pneumococcus employs a wide range of transporters to import necessary nutrients, including sugars (14), amino acids (15, 16) and essential metal ions, such as zinc (17, 18) and manganese (19, 20). Furthermore, competence, one of the hallmark characteristics of the pneumococcus, is a response to living in a diverse ecosystem and is required to relieve stress and/or to acquire beneficial genetic material from related strains and other bacteria (21, 22). Indeed, competence is induced by several stress factors, including DNA damage (23–25), and can be induced by coincubation with epithelial cells (26).

Interestingly, the pneumococcus has only a limited genetic potential to express the dedicated gene products required to survive in these new and highly varied conditions. Sequenced pneumococcal genomes from clinical and model strains reveal the presence of a pan-genome of approximately 3,000 to 5,000 genes (27), with up to 90% gene conservation between strains. Nevertheless, individual pneumococcal strains have a relatively small genome with around two million base pairs (bps). For example, strain D39V, one of the work horses of pneumococcal research, has 2,046,572 bps with 2,146 genomic features ((28), *see* https://veeninglab.com/pneumobrowse). Moreover, small bacterial genomes are more likely to be densely packed with genomic features, which puts a limit to the number of functional elements contained (29). One of the strategies used to circumvent this limitation is the encoding of moonlighting proteins, which can perform more than one function. For example, α-enolase, a major glycolytic enzyme, also binds human plasminogen, thereby combining carbon metabolism and cellular adhesion in one molecule, helping to reduce genome size (30, 31). In addition, gene regulation strategies that lead to transcriptional adaptation and nuanced levels of gene products might be the key to pneumococcal virulence and in-host survival. Unfortunately, no advanced studies comparing *S. pneumoniae* gene transcription under different environmental stressors exist, even though a detailed investigation into these gene expression patterns could provide invaluable information about its pathogenicity. A better understanding of how *S. pneumoniae* uses its compact genome to adapt to such varying environments will help guide future prevention and treatment strategies for this often deadly bacterium.

Here, we precisely quantified the transcriptome of *S. pneumoniae* (strain D39V, CP027540, (28)) under exposure to 22 different infection-relevant conditions. Next, we classified the annotated features into genes that are highly expressed across all conditions and genes demonstrating a condition-dependent, dynamic expression. Furthermore, we generated a co-expression matrix containing the correlation value of every pair of genes. We exploited the matrix to identify a new member of the competence regulon: a small hypothetical protein encoded by SPV_0391 (*briC*). Furthermore, we provide the research community with the entire compendium of normalized expression values, exhaustive fold changes and the co-expression matrix in PneumoExpress (https://veeninglab.com/pneumoexpress), a user-friendly browsable data center, enabling easy access. Finally, in PneumoExpress, we provide the research community with direct access to PneumoBrowse (https://veeninglab.com/pneumobrowse), where users can browse the genomic environment of their gene(s) of interest. The work and data presented here provide a valuable resource to the pneumococcal and microbial research community and will expand our knowledge of *S. pneumoniae* gene regulation, increasing our ability to prevent and fight infections.

## MATERIALS AND METHODS

### Culturing of *S. pneumoniae* D39V and pneumococcal transformation

*S. pneumoniae* strain D39V (available as NCTC 14078 at Public Health England, PHE) was routinely cultured without antibiotics. Strain construction and preparation of chemically-defined media (CDMs) are described in detail in the **Supplementary Materials and Methods**. Oligonucleotides are listed in **Supplementary Table S10** while bacterial strains are listed in **Supplementary Table S11**.

### Rationale for infection-relevant growth and transfer conditions of *S. pneumoniae*

The infection-relevant conditions were selected from a subset of microenvironments that the pneumococcus might encounter during its opportunistic pathogenic lifestyle. Recreating conditions that an organism naturally encounters during its lifestyle has been successful in charting a wide transcriptional responses in other bacteria (32–35). Here, we chose seven main host-like growth conditions: (i) nose-mimicking conditions (NMC), simulating colonization, (ii) lung-mimicking conditions (LMC), simulating pneumonia, (iii) blood-mimicking conditions (BMC), simulating sepsis, (iv) cerebrospinal fluid-mimicking conditions (CSFMC), simulating meningitis, (v) transmission-mimicking conditions, (vi) laboratory conditions (in C+Y medium) that allow rapid growth and (vii) co-incubation with human lung epithelial cells. Sicard’s defined medium was selected as the backbone of the first five host-like growth conditions (36). Additionally, co-incubation with epithelial cells was performed as previously described (26).

Because competence is a major hallmark of *S. pneumoniae* and it contributes to pneumococcal survival in the host (21, 22), we included three competence time-points: 3, 10 and 20 min after the exogenous addition of CSP (competence stimulating peptide-1) in C+Y. Moreover, the competence regulon is well-characterized (37, 38), allowing us to benchmark the quality of our experimental data, and analysis pipeline.

Since *S. pneumoniae* can migrate between niches, we also analyzed the transcriptomes of pneumococci being transferred between conditions. Specifically, nose to lung (NMC » LMC), nose to blood (NMC » BMC), nose to CSF (NMC » CSFMC), blood to C+Y (BMC » C+Y), C+Y to nose (C+Y » NMC), nose to transmission for 5 minutes (Transmission, 5 min), nose to transmission for 60 minutes (Transmission, 60 min), growth in nose to transmission for 5 minutes and back to nose (Transmission » NMC). Additionally, a condition mimicking meningeal fever was included, whereby *S pneumoniae* growing in CSFMC (37°C) was transferred to 40°C (FEVER). After the transfer, cells were further incubated for 5 minutes prior to RNA isolation because of the rapid production and turnover of bacterial transcripts, especially in fast-growing bacteria (39, 40). Moreover, we were interested in elucidating rapid transcriptional responses of the pneumococcus, when exposed to new and different conditions. Collectively, the 22 growth and transfer conditions are referred to as ‘infection-relevant conditions’ (**Figure 1** and **Table 1**).

**Table 1.** List of infection-relevant conditions.

| Conditions | Description | Libraries |
| --- | --- | --- |
| NMC | Growth in nose-mimicking conditions | 1,2 |
| Transmission, 5 min | Growth in NMC, transmission 5 min | 3,4 |
| Transmission, 60 min | Growth in NMC, transmission 60 min | 5,6 |
| Transmission » NMC | Growth in NMC, transmission 5 min, back to NMC 5 min | 7,8 |
| NMC » LMC | Growth in NMC, in lung-mimicking conditions (LMC) for 5 min | 9,10 |
| LMC | Growth in lung-mimicking conditions | 11,12 |
| NMC » BMC | Growth in NMC, in blood-mimicking conditions (LMC) for 5 min | 13,14 |
| BMC | Growth in blood-mimicking conditions | 15,16 |
| BMC » C+Y | Growth in BMC, in C+Y 5 min | 17,18 |
| NMC » CSFMC | Growth in NMC, in CSF-mimicking conditions (CSFMC) for 5 min | 19,20 |
| CSFMC | Growth in CSF-mimicking conditions | 21,22 |
| FEVER | Growth in CSFMC, then 40°C (fever-like) for 5 min | 23,24 |
| C+Y » NMC | Growth in C+Y, in NMC for 5 min | 25,26 |
| C+Y | Growth in C+Y | 27,28 |
| CSP, 3 min | Growth in C+Y, CSP (competence-stimulating peptide) 3 min | 29,30 |
| CSP, 10 min | Growth in C+Y, CSP 10 min | 31,32 |
| CSP, 20 min | Growth in C+Y, CSP 20 min | 33,34 |
| Infection, 0 mpi | Co-incubation with A549, 0 minutes post infection | 35,36 |
| Infection, 30 mpi | Co-incubation with A549, 30 minutes post infection | 37,38 |
| Infection, 60 mpi | Co-incubation with A549, 60 minutes post infection | 39,40 |
| Infection, 120 mpi | Co-incubation with A549, 120 minutes post infection | 41,42 |
| Infection, 240 mpi | Co-incubation with A549, 240 minutes post infection | 43 |

**Figure 1.**
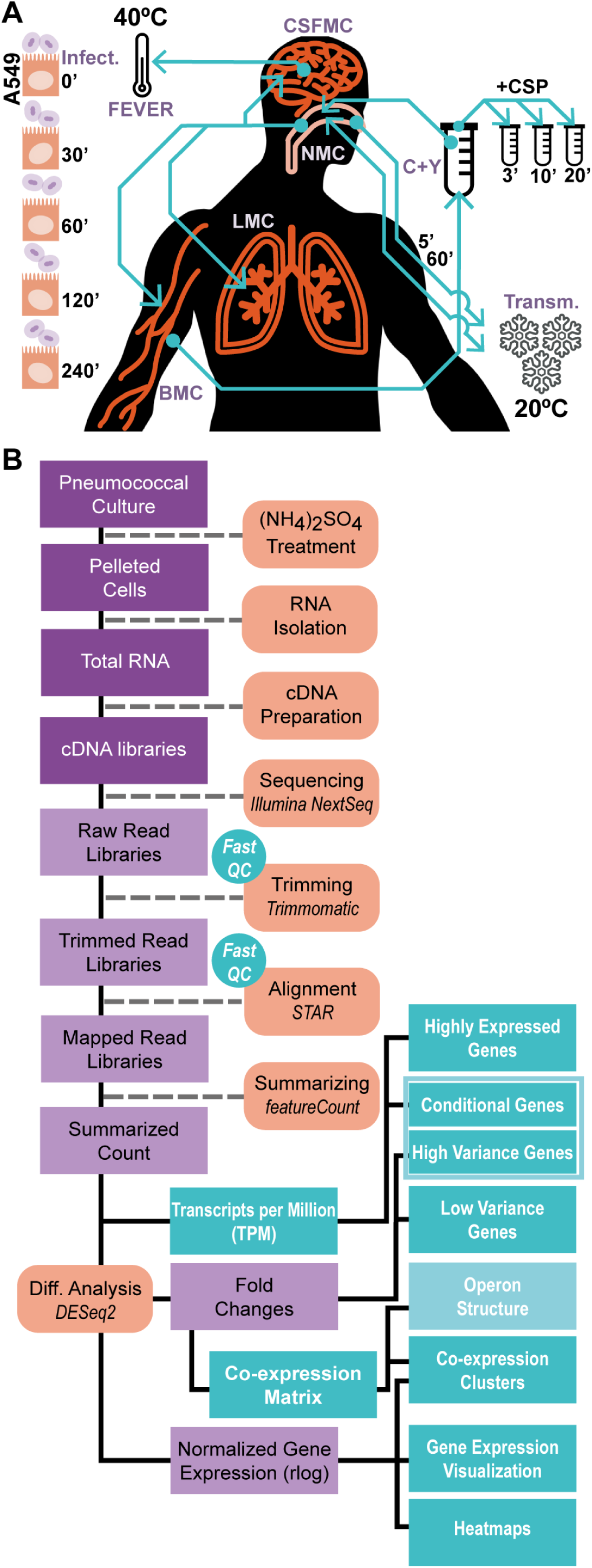
Mimicking conditions relevant to the opportunistic pathogen lifestyle. **A**. Twenty-two conditions were selected, including growth in five different conditions (laboratory, in C+Y medium [C+Y], nose-mimicking [NMC], lung-mimicking [LMC], blood-mimicking [BMC] and cerebrospinal fluid-mimicking [CSFMC]); a model of meningeal fever (FEVER); transmission conditions; eight transfers between conditions; three competence time-points and five epithelial co-incubation time-points (**Table 1**). **B**. cDNA libraries were prepared without rRNA depletion. Quality controls of the reads were performed before and after trimming. Trimmed reads were aligned and counted. Next, highly and conditionally expressed genes were categorized based on normalized read counts, while high- and low-variance genes were classified based on fold changes. High-variance and conditionally expressed genes together were defined as dynamic genes.

To recapitulate host-like growth conditions, we manipulated sugar type and concentration, protein level, partial CO_2_ pressure, temperature, acidity of medium and presence of epithelial cells (**Table 2**). We manipulated the type of carbon source because *S. pneumoniae* can utilize at least 32 different carbon sources (14), it devotes a third of all transport mechanisms to carbohydrate import (41), and it generates ATP exclusively from fermentation (42). In healthy nose and lung, respiratory mucus is the sole available carbon source, ranging from 1 g/L (lung, (43)) to 2 g/L (nasopharyngeal passage, (44)). In human mucus, Nacetylglucosamine (GlcNAc) is the main monosaccharide accounting for up to 32% of dry weight, followed by galactose (29%), sialic acid, fucose and N-acetylgalactose (45). On the other hand, glucose can be found in high concentrations in blood (46). Therefore, two sources of carbon were included: GlcNAc in NMC and LMC and glucose in BMC, CSFMC, C+Y and infection conditions.

**Table 2.** Parameters in infection-relevant growth media. *This study.

|  | Glucose<br>(g/L) | GlcNAc<br>(g/L) | Serum albumin<br>(g/L) | CO2<br>(%) | Temp.<br>(°C) | pH | Epithelial cells<br>(A549) |
| --- | --- | --- | --- | --- | --- | --- | --- |
| Nose-mimicking conditions (NMC) | - | 1.28<sup>44,45</sup> | 1<sup>44</sup> | N.D. | 30<sup>47</sup> | 7.0<sup>56</sup> | - |
| Lung-mimicking conditions (LMC) | - | 0.64<sup>43,45</sup> | 3<sup>43</sup> | 5 | 37 | 7.0<sup>56</sup> | - |
| Blood-mimicking conditions (BMC) | 0.9<sup>46</sup> | - | 67<sup>46</sup> | 5 | 37 | 7.4<sup>46</sup> | - |
| CSF-mimicking conditions (CSFMC) | 0.45<sup>57</sup> | - | 0.45<sup>57</sup> | 5 | 37 | 7.8<sup>58</sup> | - |
| Transmission | - | - | - | N.D. | 20 | - | - |
| C+Y | 1.79* | - | 0.73* | N.D. | 37 | 6.8* | - |
| Infection | 2.0<sup>26</sup> | - | 10<sup>26</sup> | 5<sup>26</sup> | 37<sup>26</sup> | 7.4<sup>26</sup> | + |

Temperature was maintained at 37°C for all conditions except for NMC (30°C, (47)) since nasal temperature ranges from 30-34°C. We set fever temperature at 40°C and transmission at 20°C (room temperature). In particular, transmission was modeled by exposing the pneumococcus to room temperature and ambient oxygen level on a sterile surface. Additionally, *S. pneumoniae* survives up to 120 minutes in transmission conditions. On the other hand, confluent epithelial cells present a biotic surface that necessitates a different pneumococcal phenotype, such as forming biofilms (48). Furthermore, the epithelial layer actively interacts with the bacteria and regulates its own transcriptome in response to the presence and activity of the pneumococcus (26). Here, we included the interaction between *S. pneumoniae* D39V and human epithelial cell line, A549 in five consecutive time points, 0, 30, 60, 120 and 240 minutes post infection (mpi, (26)). Interaction with host cells has been exploited in a similar transcriptional study in *Staphylococcus aureus* (33).

For each condition, two biological replicates were included, except for a single replicate for the last time point of the infection to A549 (Infection, 240 mpi). For a detailed description of infection-relevant conditions, *see Supplementary* **Methods**. A complete list of medium components is available as **Supplementary Table S1**.

### Total RNA isolation, library preparation and sequencing

Pneumococcal cultures from infection-relevant conditions were pre-treated with ammonium sulfate to terminate protein-dependent transcription and degradation. Total RNA was isolated and the quality checked using bleached gel (49) and chip-based capillary electrophoresis. Only co-incubation samples (libs. 35-43, **Table 1**) were depleted for rRNAs. Then, cDNA libraries were created, and sequenced on Illumina NextSeq 500 as described previously (26).

### Data analysis and categorization of genes

Quality control was performed before and after trimming. Trimmed reads were aligned to the recently sequenced genome (CP027540) and counted according to the corresponding annotation file (28). In particular, counting was performed in (i) multi-mapping mode to account for the possibility of multiple loci in the genome; (ii) in overlapping mode for genes belonging to the same operon and (iii) in fraction mode to distribute the reads coming from multimapped and overlapping reads (50). Reads were then normalized as transcripts per million (TPM, (51)) and as regularized log, a data transformation method in DESeq2 (52). Highly expressed and lowly expressed genes were categorized from TPM values excluding rRNAs. Decile values were used to partition expression values into 10 classes. The ninth decile serves as the minimum value for highly expressed genes while the first decile was used as the maximum limit for lowly expressed genes. 61 genes had TPM values above the ninth decile in all infection-relevant conditions. Along with the 12 rRNA loci, these 72 genes were categorized as highly expressed genes. On the other hand, there was no gene below the lower threshold in all conditions. However, 496 genes have TPM below the limit in at least one condition; these genes were categorized as conditionally-expressed.

Exhaustive fold changes were calculated for every pair of conditions out of the 22 infection-relevant conditions, resulting a total of 231 comparisons (**Supplementary Table S8**, (52)). Then, fold changes for comparisons reported by DESeq2 as ‘low mean normalized count’ were manually set to 0. ‘Low mean normalized count’ denotes lowly expressed genes for which significance (*p*-value) cannot be calculated confidently (52). Conditionally-expressed genes were excluded from the calculation of the limits of high and low variance genes because, by definition, these genes have high variance. The coefficient of variance (cvar) for every gene across the 231 fold changes was calculated and used as the base for variance-based partition. Here, as previously, decile values were used to partition the fold changes into 10 classes. As before, the cvar ninth decile was chosen as the minimum value for high variance genes, and the first decile as the maximum limit for low variance genes. There were 164 high variance genes, which we combined with conditionally-expressed genes and referred to as dynamic genes.

Calculations of rRNA fold changes required an alternative approach since normalization based on library size cannot be used on the highly abundant rRNAs. Instead, the expression values of the least variable 50% of all genes (1,067 features) was used to calculate the normalization factors for individual libraries. The normalization factors were applied to the whole libraries and to normalize rRNA expression values (53). Afterwards, fold changes for the rRNA-encoding genes were calculated. Hypergeometric tests to assess enrichment were performed by the built-in function, *phyper* within the R environment (v. 3.4.2).

### Generation of the co-expression matrix

The exhaustive fold changes across the pneumococcal genome were used to calculate the correlation value of every possible set of two annotated features. First, the dot-products between fold changes of the two target genes and self-dot-products of each gene were calculated. Next, the dot-products were summed: between two target genes (*a*) and self-products (*b* and *c*). The summed dot-product was referred to as non-normalized correlation value. This value was normalized by calculating the ratio between the non-normalized value (*a*) and the geometric mean of summed fold-changes (*b* and *c*). In turn, the geometric mean of summed fold changes was calculated as the square root of the multiplication product between the summed self-products. The normalized correlation value was then mapped into the matrix by the genomic positions of both genes (**Figure 6A**).

### Online compendium

The compendium can be accessed at https://veeninglab.com/pneumoexpress. The data are stored in a MySQL database as gene expression values. Gene expression graphs are generated by D3 (Data Driven Documents, https://d3js.org). Gene expression is presented in DESeq2-normalized values, rlog (52), and TPM (51) transcripts per million, including log-transformed TPM and centered TPM. Exhaustive fold changes and correlation values were included as part of the pneumococcal compendium. Centering has been described to be a useful transformation to present –omics data (54).

### Luciferin assays

Firefly luciferase (*luc*) was transcriptionally fused to the 3’-end of target operons, *comCDE* and SPV_0391-2157, to monitor gene expression levels (55). A kanamycin resistance cassette under a constitutive promoter was used as selection marker. Plate assays were performed in C+Y with 0.25 mg/ml luciferin and with and without the addition of 100 ng/µl CSP-1 from the beginning of the experiment or after 2 hours incubation. Total sugar molarity was maintained at the same level in experiment involving various sugars (*appABCD_luc*).

## RESULTS

### Infection-relevant conditions: creating the compendium

To reveal the degree of global gene regulation occurring in *S. pneumoniae* under infection-relevant conditions, we exposed strain D39V to 22 conditions that mimic aspects of the varying host environment and quantified the resulting genome-wide transcriptional responses. The conditions and growth media were chosen to recapitulate the most relevant microenvironments the pneumococcus might encounter during its opportunistic-pathogenic lifestyle in order to determine the extent to which gene expression adapts to changing environments (**Materials and Methods**, **Figure 1A** and **Supplementary Table S1**).

Statistics pertaining to the RNA-seq data were examined to determine if the coverage was sufficient to measure differential gene expression. The total number of trimmed reads per library ranged from 26 to 149 million reads (average: 89 million reads). In the non-rRNA-depleted libraries (libs. 1 to 34, **Table 1**), 99.9% of reads were mapped onto the pneumococcal genome (range: 99.8–99.9%). As expected, most reads from these non-depleted libraries aligned to the four rRNA loci of the pneumococcus. On average 95.4% of reads mapped to rRNAs, ranging from 93.4 to 97.7%, with reads mapping to tRNAs occupying on average 0.03% of total reads (0.01–0.05%, **Figure 2A**). On the other hand, of the rRNA-depleted dual RNA-seq libraries (libs. 35 to 43, **Table 1**), an average of 64.6% of trimmed reads mapped onto the pneumococcal genome (range: 48.4%–84.6%) while the rest mapped to the human genome. Excluding reads that mapped to pneumococcal rRNA genes and taking into account the read length (75 nt), the sequencing depth of libraries (i.e. coverage of the genome) ranged from 76× to 1944×, which is sufficient to elucidate differential gene expression in bacteria (59, 60).

**Figure 2.**
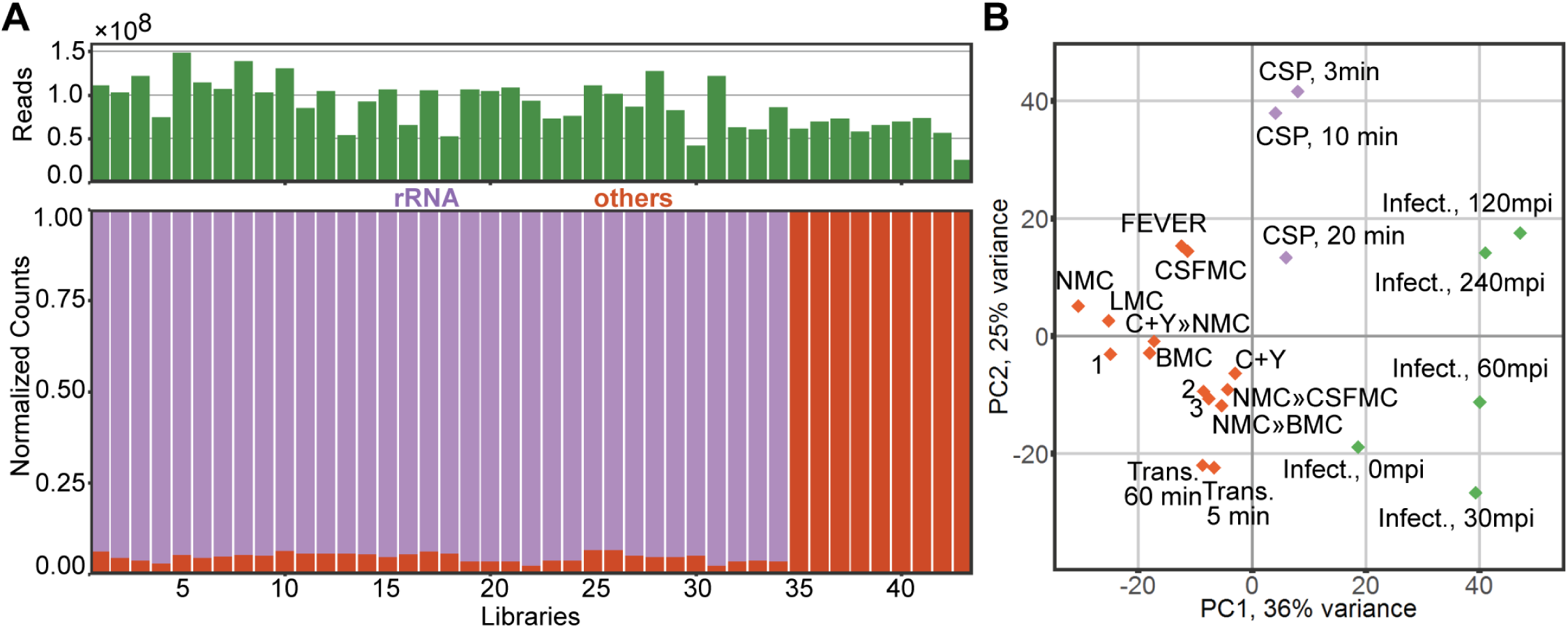
Distribution of libraries and conditions. **A**. The number of trimmed reads of the 43 libraries ranged from 26 to 149 million reads, averaging 89 million reads. Non-rRNA-depleted libraries were dominated by reads mapped to ribosomal RNA genes, averaging 95% (range: 93–98%, libs. 1–34). **B**. Principal component analysis of gene expression in all conditions showed three clusters of conditions: conditions based on competence (CSP, 3 min, 10 min and 20 min; purple), epithelial infection (infection 0 mpi, 30 mpi, 60 mpi, 120 mpi and 240 mpi; green) and other infection-relevant conditions (orange). 1 = Transmission » NMC; 2 = NMC » LMC; 3 = BMC » C+Y.

We then used principal component analysis (PCA) to describe the general behavior in all conditions, especially relating to how each condition resembled another condition, relying on the fact that close distance or clustering in a PCA plot indicates a similar transcriptomic response between these conditions. This analysis indicated that three clusters of conditions could be observed, which roughly corresponded to the basal medium used in the conditions. The first cluster consisted of the five time-points that occur during the infection of human epithelial cells, while the second cluster consisted of the competence time-points. Finally, the third cluster contained all other conditions (**Figure 2B**). Interestingly, the growth in C+Y clustered with the latter group and not with the competence samples, indicating that clustering represents the biological response and is not solely dependent on the type of medium. The clustering behavior in the PCA plot also indicated that the activation of competence in the pneumococcus represents a major transcriptional shift.

Because the dataset is derived from two different preparation and sequencing batches, we wanted to ensure that the clustering was genuine and not due to variations in sample processing. In particular, libs. 1-34 belong to a different batch than libs. 35-43 (*see* **Table 1**). In order to account for these batches, we performed batch effect correction (61). We did not observe appreciable differences in genome-wide expression values and clustering behavior before and after batch effect removal. Thus, we concluded the clustering behavior is truly due to biological responses and not because of any batch effect. Subsequently, we used the original dataset for downstream analysis.

To visualize gene expression across the 22 conditions, we generated the ‘shortest tour’ through the PCA plot, which helps to visualize which samples show the most similar transcriptomes (**Supplementary Figure S1**). First, we calculated the Euclidean distances, or the straight line, between the conditions on the two-dimensional PCA plot. Subsequently, we minimized the total distance needed to visit all the points only once, the so-called TSP algorithm (travelling salesman problem, arXiv: https://arxiv.org/abs/cs/0107034). We have further validated gene expression values by qPCR (**Supplementary Figure S2**). Taken together, we observed a large *S. pneumoniae* differential gene expression that was highly dependent on its environment.

### Categorization of genes: highly expressed and dynamic genes

Read count normalization was performed in two ways: transcripts per million (TPM, (51)) and regularized logarithm (rlog, (52)). While TPM-normalization corrects for the size of the library and length of a feature, rlog scales abundance directly to a log_2_-scale while adjusting for library size. In addition, the rlog is considered suitable for visualizing gene expression across diverse conditions (52), while TPM values were used to categorize genes as highly or lowly expressed.

The 72 highly expressed genes include those encoding rRNAs and the 34 genes coding for ribosomal structural proteins. Other genes, including the two translation elongation factors *fusA* and *tuf*, DNA-dependent RNA polymerase *rpoA*, transcription termination protein *nusB*, and the histone-like protein *hlpA*, were also highly expressed in all conditions. Additionally, a set of genes associated with carbohydrate metabolism were highly expressed: *fba* (fructose-bisphosphate aldolase), *eno* (enolase), *ldh* (lactate dehydrogenase), *gap* (glyceraldehyde3phosphate dehydrogenase), and a gene encoding a subunit of ATP synthase, *atpF*. A complete list of highly expressed genes is available as **Supplementary Table S2.** The highly expressed genes are enriched for genes with essential cellular functions (hypergeometric test, *p* < 0.05, (64, 65)), with little differential expression across conditions. We speculated that because these genes perform core housekeeping functions, such as protein translation, RNA transcription, DNA maintenance and carbon metabolism, the expression of these genes is maintained at a high abundancy with little to no transcriptional regulation under different conditions.

On the other hand, our analysis reported 498 conditionally expressed genes, which were transcriptionally regulated as demonstrated by the great fluctuation of mRNA across infection-relevant conditions. A full list of conditionally expressed genes is available as **Supplementary Table S3.** In this category, 48 genes (9.6%) encoded proteins involved in carbohydrate import, including transporters of galactosamine (*gadVWEF*), cellobiose (SPV_0232-4, *celBCD*), hyaluronate-derived oligosaccharides (SPV_0293, SPV_0295-7), galactose (SPV_0559-61), ascorbic acid (*ulaABC*) and mannose (SPV_1989-92). Out of the 48 genes, 31 genes are preceded by a catabolite control protein A (CcpA) binding site, which suggests that their expression is under the direct control of CcpA. In *S. pneumoniae*, CcpA acts as central switch that regulates carbon metabolism and contributes to pneumococcal survival and virulence inside the host (64, 65).

In our dataset, the 31 genes encoding for sugar importers were highly expressed in the presence of the alternative sugar, N-acetylglucosamine, found in the NMC and LMC media and in transfers to NMC. Indeed, the transfer from NMC (carbon source: N-acetylglucosamine) to LMC (carbon source: N-acetylglucosamine) did not lead to the differential expression of these genes. On the other hand, genes encoding for importers of cellobiose (SPV_0232-4, *celBCD*), galactose (*gatABC*), lactose (*lacE2F2*) and multiple sugars (SPV_1583-5) were activated in co-incubation with epithelial cells, although the medium only contains glucose. This activation might be due to the presence of host-derived alternative sugars, as we previously showed that washed epithelial cells did not incite such gene activation (26).

In addition, exhaustive comparisons (231 in total) between every set of two conditions were performed. The coefficient of variation of the summarized fold changes per gene were used to categorize high and low variance genes (**Materials and Methods**). High variance genes include pyrimidine-related genes (*pyrFE, pyrKDb, uraA, pyrRB-carAB*) and purine-associated genes (*purC, purM, purH*). These genes were activated during co-incubation (infection, 0 mpi to 240 mpi), transfer to transmission (transmission, 5 min and 60 min) and growth in LMC. Furthermore, we observed that members of the ComE regulon, designated as early competence genes (66), were heavily upregulated in all competence time points (CSP, 3 min, 10 min and 20 min), CSFMC, FEVER and late co-incubation with epithelial cells (infection, 120 mpi and 240 mpi). In contrast, the expression of the ComX-regulated, i.e. late competence (37, 38), genes peaks 10 minutes after the addition of CSP1 (CSP, 10 min) and on transfer to transmission (transmission, 5 min and 60 min). We have combined conditionally expressed genes and high variance genes into a single category: dynamically expressed genes (**Figure 3** and **Supplementary Table S4**). In addition, the expression of low-variance genes can be observed in **Supplementary Figure S2A**. Together, this coarse-grained analysis showed the presence of a large set of genes that are conditionally expressed (approximately 25% of all genetic features), indicating a large-scale rewiring of the pneumococcal transcriptome upon changing conditions.

**Figure 3.**
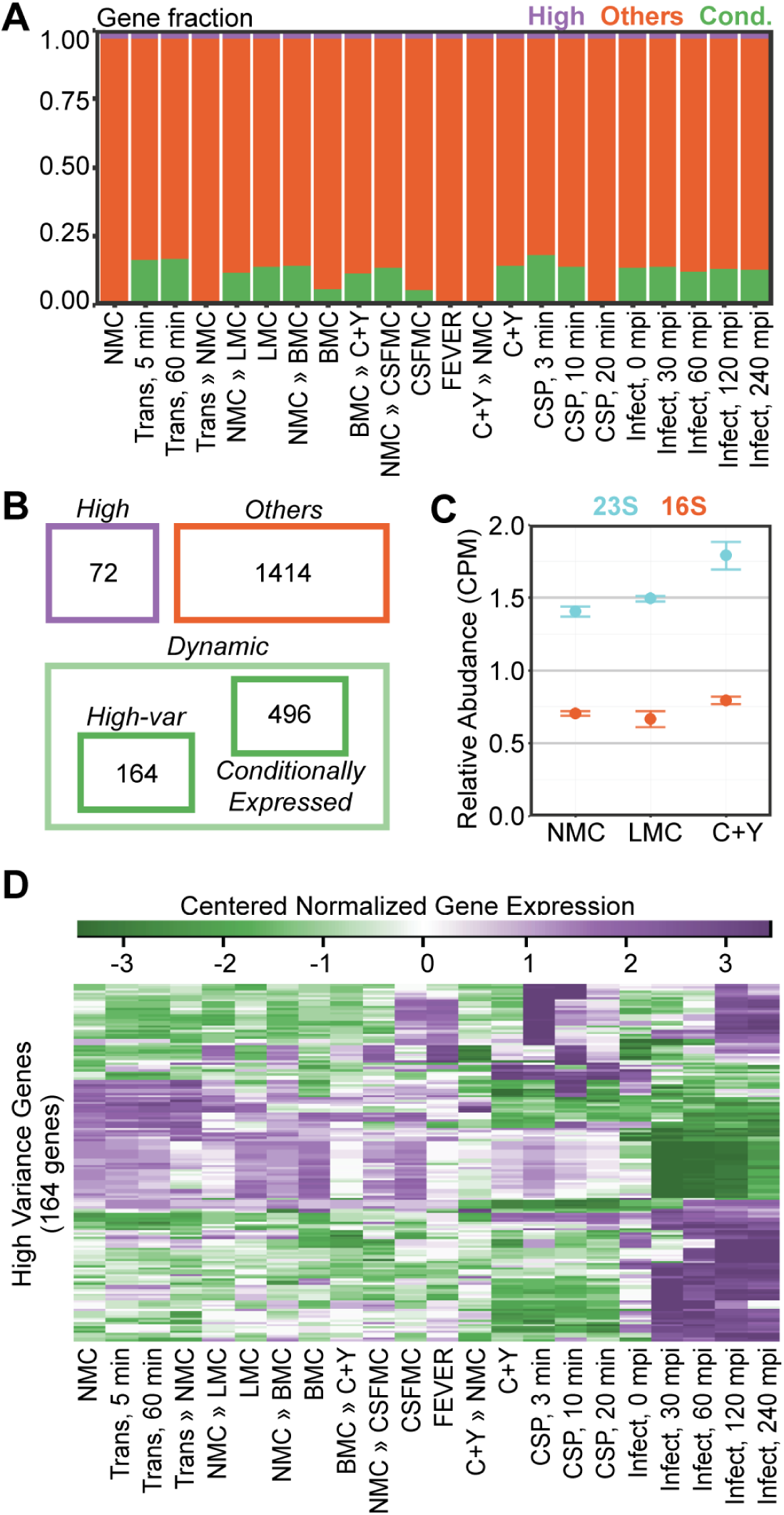
Categorization of genes. **A**. Visualization of the number of genes in all conditions according to their categories: steadily highly expressed (purple), conditionally expressed (green) and others (orange). Of the 2,146 features, 73 are classified as highly expressed, while 498 features are conditionally expressed (lowly expressed in at least one condition). **B**. Highly expressed genes include essential genes, genes encoding ribosomal proteins and rRNAs. Dynamic genes are a combination of the 165 high-variance genes and 498 conditionally expressed genes. **C**. The 23S rRNA was significantly downregulated in nose-mimicking (NMC) and lung-mimicking (LMC) growth compared to rich C+Y growth (p < 0.05). The 16S rRNA showed a similar trend, but it was not statistically significant (p = 0.33, C+Y to NMC; p = 0.83, C+Y to LMC; error bars represent standard error). **D**. Expression values (regularized log) of high-variance genes were centered, as described in **Supplementary Materials and Methods**, and plotted as heat maps. Distinct clusters of gene expression can readily be observed (purple: high expression, green: low expression).

### Growth-dependent expression of rRNA

While rRNA depletion allows for a higher coverage of mRNA sequence reads, it also introduces bias to sequenced libraries, partially due to depletion of sequences similar to rRNA (67). We have opted not to deplete rRNAs in most of the libraries, endowing the compendium with an unbiased quantification of the total RNA. This approach also gave us the rare opportunity to investigate the expression levels of ribosomal RNAs (rRNAs) in the conditions under study. Because of the abundance and stability of rRNA, we adopted an alternative normalization procedure prior to calculating the fold-change. Rather than normalizing rRNA read counts based on the total number of reads in the library (as is the standard procedure), we exploited read counts of low variance genes to define an alternative normalization factor (**Materials and Methods**). Furthermore, this approach more directly implements the basic assumption of differential gene analysis, i.e. that the majority of genes are not differentially expressed when comparing two conditions. Generally, this assumption implies that total library size is largely insensitive to differential expression of a small fraction of genes and therefore constitutes a simple and suitable normalization factor of gene expression. However, since rRNAs dominate the total RNA libraries, differential rRNA expression would have a non-negligible effect on total library size, necessitating a more direct normalization method, as applied here.

When comparing rRNA levels between the various conditions, they were significantly higher in fast-growing pneumococci (C+Y, CEP) compared to slow-growing cells (nose-mimicking and lung-mimicking growth, **Figure 3C**). rRNA expression in the Gram-positive model organism *Bacillus subtilis* has previously been reported to be regulated by availability of dGTP because the initiating nucleotide for rRNA transcription is a GTP rather than the more common ATP (68). Even though rRNA operons in *S. pneumoniae* are also initiated with GTP (28), we did not observe a correlation between the initiating nucleotide and gene expression levels in cells grown in different media (**Supplementary Figure S2C**). Nevertheless, in prokaryotes including *S. pneumoniae*, genes encoding rRNAs and proteins are conserved in a location close to the origin of replication (69–71). The *ori*-proximal location of the four pneumococcal rRNA loci results in a higher gene copy number of rRNAs in fast-growing cells, such as in C+Y, as a direct consequence of the increase in replication initiation frequency. Indeed, we find that in general, constitutively expressed genes located close to the origin of replication demonstrate higher expression under fast growth (24, 69).

### Condition-responsive expression of pneumococcal genes

Next, we clustered genes based on TPM-normalized expression values (51, 72) in order to identify relevant clusters that would indicate condition-responsive expression. Doing this, we recovered a 19-gene cluster that was responsive to the temperature increase between transfers. Temperature is one of the major physicochemical properties in our model, ranging from room temperature, 30°C, 37°C and 40°C across infection-relevant conditions. For example, we observed a high fold change of the cluster compared to other genes when comparing CSFMC to FEVER, representing the shift from 37°C to 40°C (**Figure 4A**). We also noticed a similar fold change when comparing NMC growing at 30°C to LMC, BMC and CSF-MC growing at 37°C. In the comparison between NMC (at 30°C) to transmission at 20°C (trans ***»*** NMC), we did not detect appreciable upregulation of this cluster. The absence of upregulation can also be observed in the comparison between C+Y (37°C) to C+Y » NMC (30°C) and in comparison, between BMC (37°C) to BMC » C+Y (37°C). Members of the heat-responsive cluster include genes encoding well-known heat-shock proteins, *hrcA-grpE-dnaK-dnaJ* and *groESL,* and genes encoding components of the proteolytic Clp complex, *clpL, clpP* and *clpC*. Other heat-responsive members include *ctsR* (in the same operon as *clpC*), SPV_2171, (in the same operon as *hrcA*), *glnQ2* (encoding for glutamine ABC transporter), *cbpA* (encoding choline binding protein A), SPV_2019-20 (pneumococcal two component system) and *thiXYZ*-SPV_2027 (thiamin ABC transporter).

**Figure 4.**
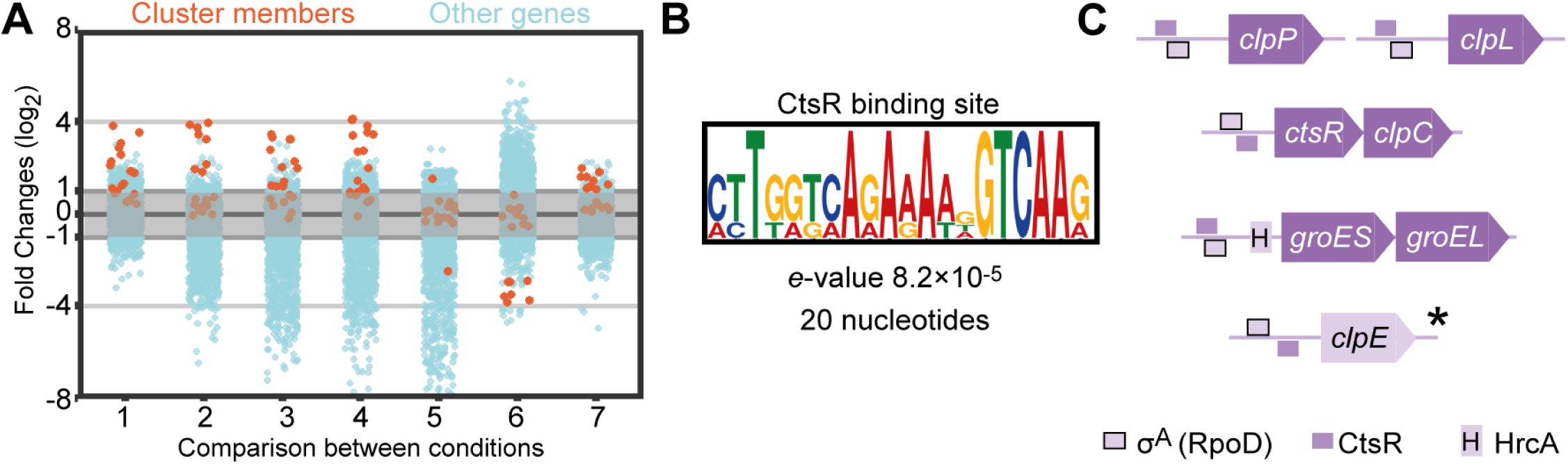
Temperature-responsive genes. **A**. A cluster containing 19 genes was recovered from the clustering analysis based on the TPM values across 22 conditions. Comparisons were selected that demonstrate the fold change of cluster members (log2) in orange as compared to those of all other genes in light blue. Comparison 1 is CSFMC to FEVER; 2 is NMC to NMC » LMC; 3 is NMC to NMC » BMC; 4 is NMC to NMC » CSFMC; 5 is NMC to Trans, 5 min; 6 is C+Y to C+Y » NMC; and 7 is BMC to BMC » C+Y. **B**. Motif enrichment analysis between 60 and 10 nucleotides upstream of the transcription start sites of the six-membered operons resulted in a 20-nt wide CtsR binding site, CTTGACHTTTTCTGACCAAG. **C**. A genome-wide search for CtsR sites recovered four operons with a reported CtsR site that belonged to the original 19-gene cluster and one other gene, *clpE*. CtsR sites overlap with RpoD sites, and *groESL* expression is co-regulated by HrcA. \**clpE* is preceded by non-overlapping RpoD and CtsR binding sites.

Next, we performed motif enrichment analysis on the upstream regions of the abovementioned operons and recovered the CtsR binding site, which preceded five operons across the genome (**Figure 4B**). This common binding site for a regulator of heat-shock genes indicates that the expression levels of these genes are all controlled by the same transcriptional signals, meaning that they are highly interrelated. The identified motif is very similar to the predicted orthologous motif in closely related species except for the strong presence of one thymine (T) preceding the reported motif (73). Four of the five operons, *clpP*, *clpL*, *ctsR-clpC* and *groESL* belong to the 19 gene cluster of temperature-responsive genes, which also contains the HrcA regulon. Finally, the monocistronic fifth operon is constituted by *clpE* and also has a CtsR binding site. However, it did not cluster with the heat-responsive genes (**Figure 4C**) and has a rather stable constitutive expression across the conditions. While the promoter regions of all five operons contain recognition sites for both CtsR and RpoD (***σ***^A^), the primary sigma factor during growth, we speculate that the position of these sites relative to each other determines the efficiency of CtsR control. Indeed, these sites do not overlap in the promoter of *clpE* (*see Figure* **4C**), thereby potentially limiting the effect size of CtsR-mediated repression and derepression. Additionally, the *groESL* operon contains an HrcA binding site in addition to the RpoD and CtsR sites. Together, these analyses show that, aside from highly-conserved heat-shock proteins, *S. pneumoniae* D39V activates specific genes in elevated temperature, including those encoding for virulence factor CbpA and proteins with as of yet unknown function, SPV_2027 and SPV_2171.

Additionally, we analyzed genome-wide expression values using the TPM to discover genes that were upregulated in only one condition. A functional enrichment analysis (hypergeometric test) revealed that several ABC transporters were enriched in the subsets of condition-specific genes. The expression of genes encoding two sugar transporter complexes, *malXCD,* transporting maltose/maltodextrin, and *msmEFG*, transporting multiple sugars, are specific to one condition. While *malXCD* is strongly upregulated 30 minutes after the infection of A549 cells, *msmEFG* is activated most strongly in NMC and to some degree in LMC and C+Y ***»*** NMC (**Figure 5A**). The strong activation of *malXCD* can be attributed to the presence of host-derived sugars (26). On the other hand, *msmEFG*, along with *agaN* (encoding ***α-galactosidase) is under the control of*** CcpA, which in turn is regulated by the intracellular levels of glucose found in low levels in the nose-mimicking and lung-mimicking mediums. Furthermore, the expression of genes encoding the branched-chain amino acid (BCAA) transporter, *livFGMHJ*, is activated in infection conditions at 30 and 60 mpi, and this expression can be partly explained by the presence of a CodY binding site and the presence of host-derived leucine, isoleucine and valine. Interestingly, an oligopeptide ABC transporter, *appABCD*, is activated upon the transfer between C+Y to NMC (C+Y » NMC). While the level of bovine serum albumin as source of oligopeptides in the two conditions is comparable (0.73 g/L in C+Y and 1 g/L in NMC), the major difference between these two conditions is the carbon source, glucose in C+Y and N-acetylglucosamine in NMC. A closer inspection of the immediate upstream region of *appABCD* revealed that aside from an RpoD site, the operon is also preceded by a CcpA site (**Figure 5B**). To investigate *appABCD* expression and its response to different carbon sources, we transcriptionally fused firefly luciferase (*luc*) behind *appD*. Three different carbon source compositions were studied in a C+Y background: glucose, N-acetylglucosamine and an equimolar combination of glucose and Nacetylglucosamine. Growth was slightly affected by changes in the carbon source, and *appABCD* expression was varied – repressed in the presence of glucose and upregulated in increasing levels of Nacetylglucosamine, indicating that CcpA regulates the transcription of this oligopeptide transporter. Indeed, a previous array-based study also suggested the operon to be part of the CcpA regulon (64).

**Figure 5.**
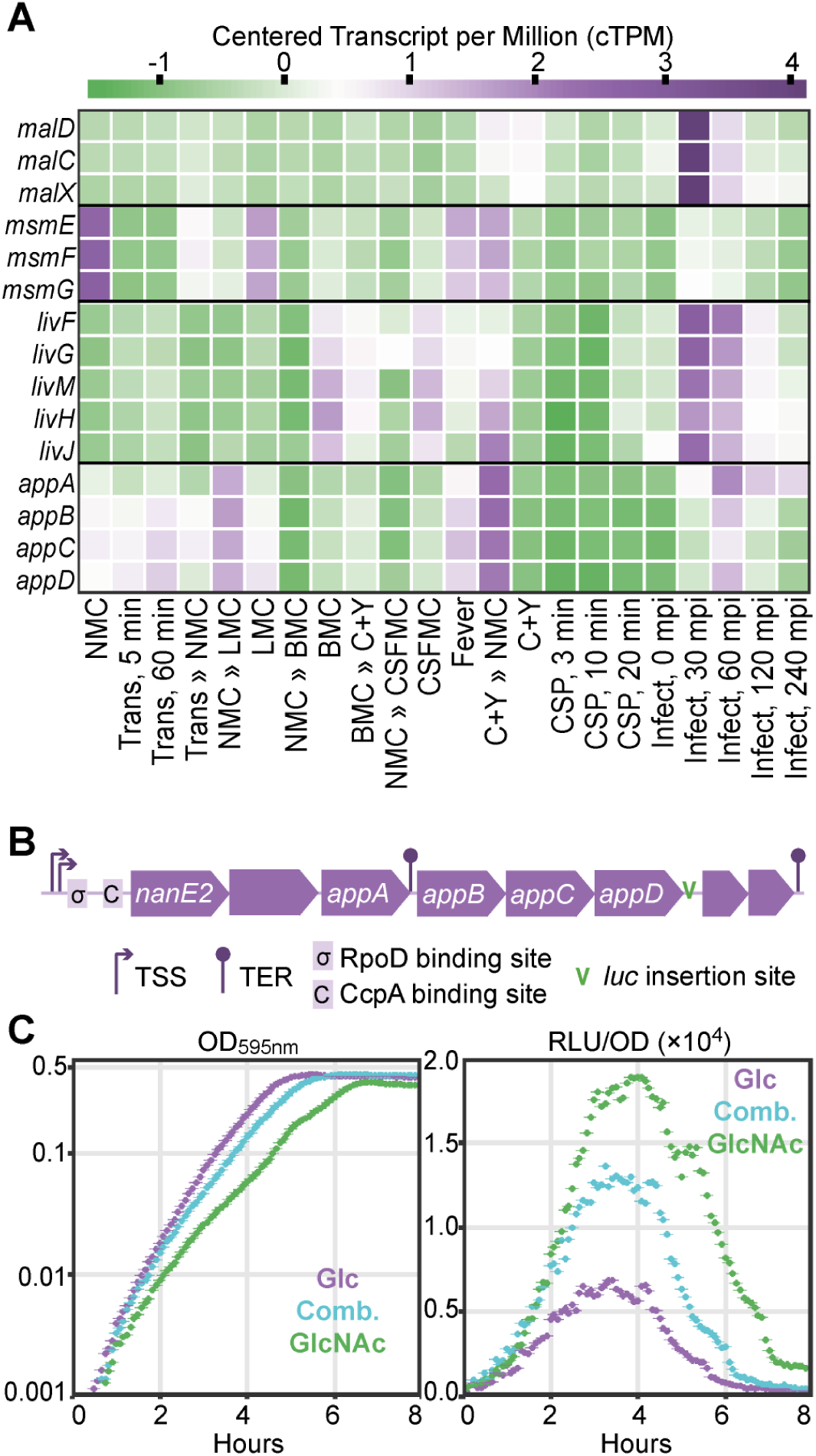
Condition-specific gene expression. **A**. ATP-binding cassette (ABC) transporters are strongly overrepresented among condition-specific genes. The expression of *malXCD*, which encodes the maltose/maltodextrin transporter, peaks in Infection, 30 mpi while *msmEFG*, which encodes a multi-sugar transporter, is highly expressed in NMC and to a lesser degree in LMC, C+Y » NMC and FEVER. Infection conditions (30 and 60 mpi) incite the expression of *livFGMHJ*, which encodes the branched-chain amino acid transporter, and the transfer between C+Y to NMC (C+Y » NMC) activates the expression of *appABCD*, encoding an oligopeptide transporter. Purple indicates high expression and green indicates low expression, as indicated by the legend above the graph. **B**. The upstream region of *appABCD* contains the RpoD and CcpA binding sites. Firefly luciferase (*luc*) is transcriptionally fused after *appD*. **C**. While growth is barely affected by different carbon sources, the luciferin signal increases in the presence of N-acetylglucosamine. Glc: glucose; Comb.: equimolar combination of glucose and N-acetylglucosamine; GlcNAc: N-acetylglucosamine.

### Assembly of genome-wide correlation values to generate a co-expression matrix

To facilitate the identification of operon structure and regulons, we created a co-expression matrix based on the fold changes in expression levels between the conditions. First, we exhaustively compared genome-wide fold changes between every two conditions of the 22 infection-relevant conditions. Next, we calculated the dot-product of the vectors containing all the fold changes of gene 1 with the vector containing all the fold changes of gene 2 (*a*, non-normalized correlation value). Similarly, we determined the self-dot-products of gene 1 (*b*) and gene 2 (*c*). A normalized correlation value was obtained by calculating the ratio of the non-normalized value (*a*) to the geometric mean of the self-dot-products (*b* and *c*). We then mapped this correlation value according to the genomic positions of the original genes (**Figure 6A** and **Materials and Methods**). Previously, a similar method was exploited to generate a co-expression matrix across different eukaryotic species to recover genetic modules (74). The maximum correlation value, including self-correlation, is 1, and the determined correlation values range from −0.97 to 1.

**Figure 6.**
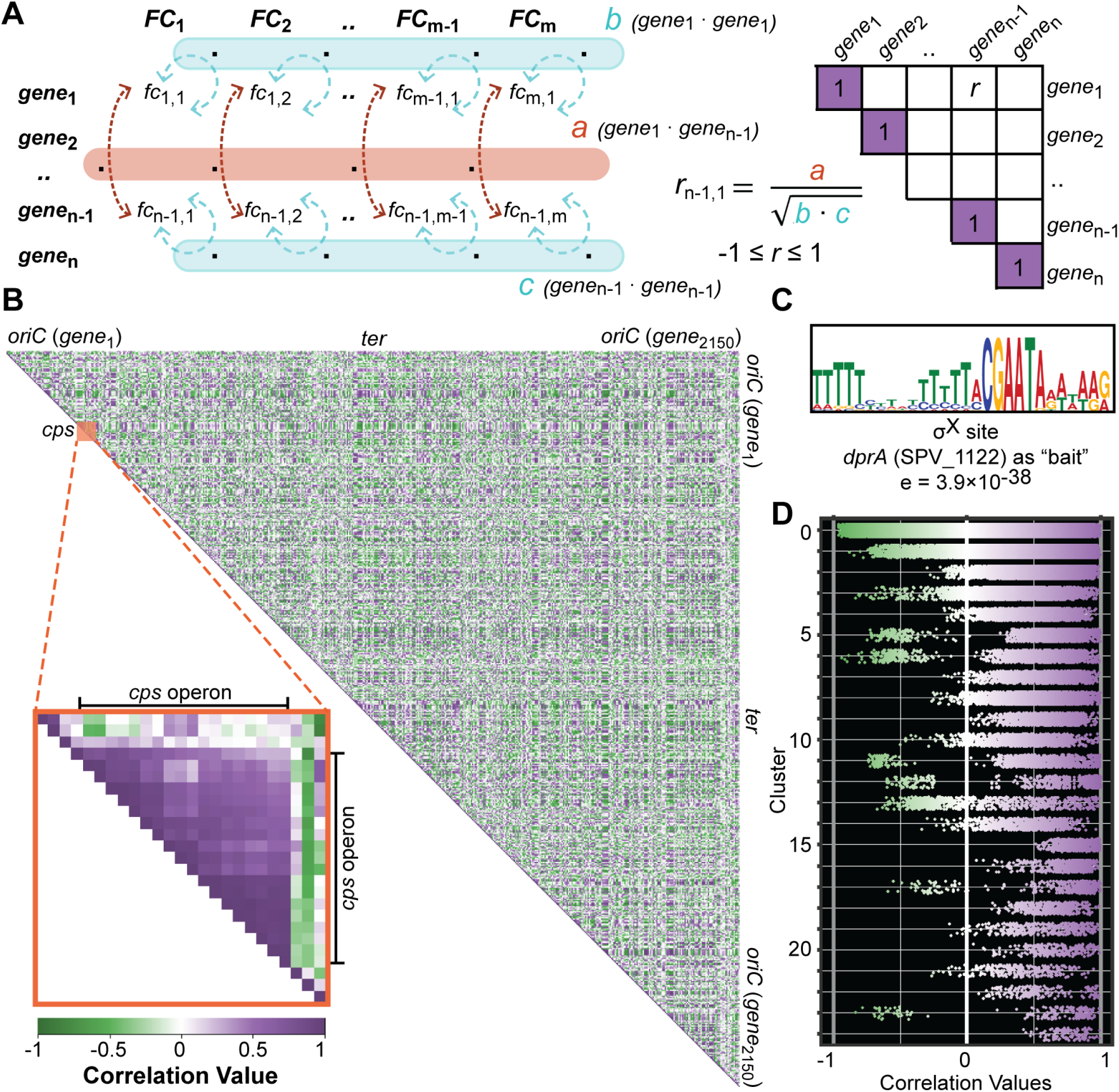
Assembly of the co-expression matrix from the correlation values of every two pneumococcal genes. **A**. The exhaustive fold change calculated for every set of two genes are converted into a correlation value: first, the dot-product between two genes (*a*, orange) and the dot product of each gene with itself (*b* and *c*, blue) are calculated. The correlation value is the ratio between a and the geometric mean of *b* and *c*. Values were assembled by the genomic coordinates of the target genes. **B**. The co-expression matrix as a visualized gene network. Self-correlation values are 1 by definition and correlation values were plotted according to the genomic positions of target genes. Purple and green indicate positive and negative correlation values between two genes, respectively. Color intensities represent correlation strength. Blocks of highly correlated genes close to the matrix diagonal indicate operon structures, for example for the *cps* operon (inset). **C**. An enriched promoter motif recovered from genes highly correlated to *dprA* (SPV_1122) matches the consensus ComX binding site (76). **D**. Pneumococcal genes were clustered into 25 clusters based on TPM (transcripts per million). Then, correlation values for every two genes within each cluster were plotted. Cluster 0 is non-modular, and its correlation values can be considered as random. Within-cluster values showed a clear trend towards higher correlation (purple).

Around the matrix diagonal, we observed blocks of highly correlated genes, indicating their co-expression and proximity. These proximity blocks are referred to as ‘putative operons’ and are used as input for further analysis (28). In particular, the well-known *cps* operon (75) can be observed in the co-expression matrix in which 16 consecutive genes are co-expressed as a single operon (**Figure 6B** inset). In contrast, the correlation values between members of the *cps* operon and genes either upstream or downstream of the locus are considerably lower.

In addition to belonging to the same operon, co-expression can be mediated by shared expression-regulatory properties. Regulatory proteins typically interact with the promoter regions of regulated genes and identifying groups of genes that are regulated by the same regulatory protein (or RNA) is of particular interest in the characterization of the pneumococcal response to a changing environment. From the matrix, we recovered 46 features (of 26 operons) that are highly correlated to *dprA,* a member of the ComX regulon. Motif-enrichment analysis on the 50-nt region upstream of the corresponding 26 start sites resulted in a 28 nucleotide motif (**Figure 6C**) that closely matched the ComX binding site as previously reported (76). Furthermore, we clustered pneumococcal genes based on their normalized expression value (TPM) and recovered 25 clusters (72, 77). The first cluster, cluster 0, is a non-modular cluster that contains all the genes that did not fit into any of the other clusters. This cluster can therefore be considered as a random control. When we plotted the correlation values of every set of two genes within each cluster, we observed a bias towards higher correlation values in all clusters except for the non-modular cluster (**Figure 6D**). As an additional control, we selected 120 random genes divided into 3 groups and plotted the correlation values within the groups. There, we observed a truly random distribution of correlation values in all groups (**Supplementary Figure S3A**).

Finally, we hypothesized that a pair of genes with strong correlated expression across infection-relevant conditions is likely to share a cellular function. We concluded that the co-expression matrix represents a simple network of genome-wide expression profiles that reveal meaningful transcriptomic responses to a changing environment. Moreover, by comparing gene expression profiles across a wide range of conditions, direct and indirect regulatory connections between genes can be unveiled. On the other hand, the co-expression matrix also has the potential to elucidate negative regulators indicated by strong negative correlation values with their target genes. Unlike previous reports (74, 78), the coexpression matrix that we describe here does not decompose pneumococcal genes into clusters of co-expressed genes. Rather, by extracting correlation values between a gene against all pneumococcal genes, we can ‘fish’ for co-expressed genes to generate starting hypotheses and further assist in the design of downstream experiments to elucidate the cellular function of hypothetical gene(s).

### Exploiting the matrix to reveal a new member of the competence regulon

Two-component regulatory systems (TCSs), consisting of a sensor histidine kinase that senses an environmental stimulus, and a cognate response regulator that controls gene expression after activation by the kinase, are essential for adapting to the microenvironment and fine-tuning gene expression in the pneumococcus (79, 80). ComDE, the best-described TCS, is controlled by a quorum-sensing mechanism and regulates competence, or X-state, which in turn is responsible for the expression of ∼100 genes and a wide range of phenotypic changes (80, 81). By extracting correlation values of all pneumococcal genes, we recovered genes strongly correlated with *comE* that encoded the TCS DNAbinding response regulator. Specifically, we identified 26 *comE*-associated genes with correlation values above 0.8. ComE autoregulates its own expression along with the expression of *comC1* (SPV_2065) and *comD* (SPV_2064), which belong to the same operon and indeed correlate strongly with *comE*. Furthermore, other known members of the ComE regulon, such as *comAB* (SPV_0049-50), *comW* (SPV_0023) and *comM* (SPV_1744) belong to the same cluster.

Interestingly, SPV_0391, encoding a conserved hypothetical protein, was included in the group. SPV_0391 has not been previously reported as part of the competence regulon in array-based pneumococcal competence studies (37, 38). Furthermore, *comE*-associated genes are not localized in a specific genomic location, but are spread out throughout the genome (**Figure 7A**), ruling out the effect of genomic location. Expression values of *comCDE* and SPV_0391 across infection-relevant conditions demonstrated a strong correlation between the genes (**Figure 7B**). In the promoter region of SPV_0391, we observed a ComE-binding site consisting of two ComE-boxes, which suggests direct regulation by ComE. To study the expression of SPV_0391 and the responsiveness of the identified ComE site, we transcriptionally inserted firefly luciferase (*luc*) downstream of SPV_0391, immediately followed by the pseudogene *ydiL* (SPV_2157), which contains a frameshift mutation after position 165 and is unlikely to be translated into a functional protein. Importantly, no terminators or additional transcription start sites were detected between SPV_0391 and *ydiL*, suggesting they form an operon together. The presence of a small hypothetical protein, SPD_0392, was previously annotated within the *ydiL* coding region, so we chose to integrate *luc* downstream of SPV_0392 to avoid potential downstream effects (**Figure 7C**). We compared the luminescence signal in this strain to that of a D39V derivative that expressed *luc* transcriptionally fused to the 3’-end of *comCDE* (24).

**Figure 7.**
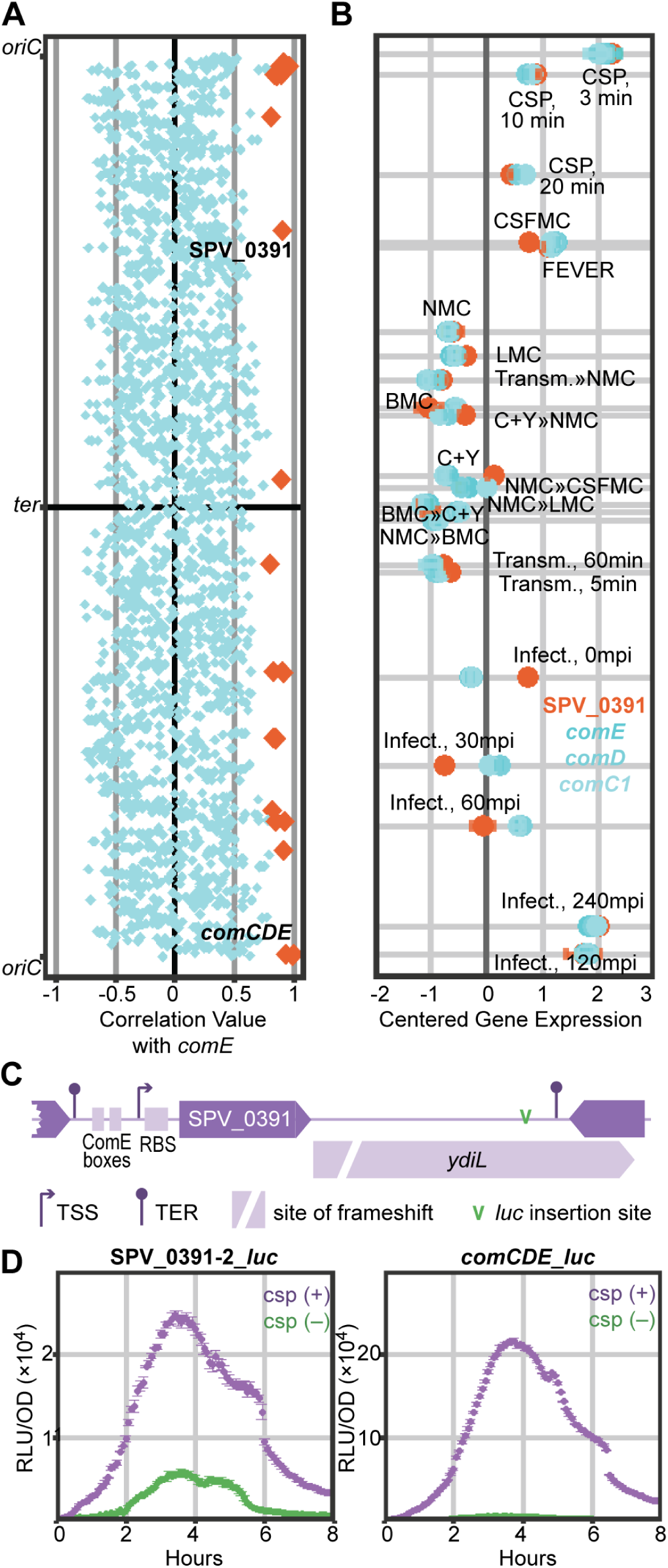
The co-expression matrix reveals a new competence-regulated gene. **A**. The gene encoding the pneumococcal response regulator, ComE, was used to recover 26 highly correlated features (orange diamonds). The group is mainly populated by known members of the ComE regulon, except for SPV_0391, a conserved hypothetical gene not previously reported to be part of the competence regulon. **B**. Centered regularized log as expression values of SPV_0391 (orange) and *comCDE* (shades of blue) were plotted against the shortest tour of infection-relevant conditions. Expression values of the four genes closely clustered together. **C**. Genomic environment of SPV_0391 with two preceding ComE boxes. SPV_0391 shared an operon structure with a pseudogene, *ydiL*. **D**. Firefly luciferase (*luc*) was transcriptionally fused downstream of SPV_0391 or *comCDE* to characterize their expression profiles with and without the addition of exogenous competence stimulating peptide-1 (CSP-1, 100 ng/µl). The addition of exogenous CSP-1 incited similar luminescence profiles in SPV_0391-SPV_0392-*luc* and in *comCDE*-*luc* strains.

Exogenous addition of 100 ng/µl of CSP-1, to stimulate competence, stimulated luciferase activity in both reporter strains (**Figure 7C, Supplementary Figure S3B**). Although the luciferase signal from SPV_0391 was an order of magnitude lower than the luminescence driven by *comCDE,* the signal profiles were very similar. The difference in signal strength may stem from a weaker promoter driving SPV_0391 than *comCDE*.

We disrupted SPV_0391 to elucidate its role in pneumococcal competence and found that deletion of this conserved feature did not affect growth in C+Y or the expression profiles of luciferase downstream of *comCDE* and *ssbB*, members of the ComX regulon (*not shown*). Finally, transformation efficiency in the deletion strain was not significantly different from that of the parental strain. Thus, while SPV_0391 is under the control of ComE and part of the pneumococcal competence regulon, we could not determine its role in pneumococcal competence. Indeed, recent work has shown that the protein encoded by SPV_0391 (named BriC) does not play a role in transformation, but rather promotes biofilm formation and nasopharyngeal colonization (bioRxiv: https://doi.org/10.1101/245902). The fact that we could identify SPV_0391 (*briC*) as a bona fide member of the ComE regulon, though array-based technology could not, demonstrates the advantages of RNA-seq over hybridization technology (37, 38).

### Development of an interactive data center to explore gene expression and correlation

To enable users to easily mine the rich data produced here, we developed an interactive data center accessible from https://veeninglab.com/pneumoexpress where users can easily extract expression values and fold changes of a gene of interest, as well as quantitative information on how its expression profile correlates with that of other genomic features (**Figure 8**). As a proof of principle, in addition to the competence regulon, we demonstrated results obtained by examining the PyrR regulon. Traditional transcription factors bind to the promoter region of a DNA molecule and the confident prediction of all their binding sites is challenging. PyrR, on the other hand, controls the expression of its regulon through an interaction with an RNA switch (82, 83). We identified four of these RNA switches (in front of *uraA*, *pyrFE*, *pyrRB*-*carAB* and *pyrKDb*) that are predicted to regulate the expression of nine genes based on putative operon structures (28). As expected, the eight genes showed a strong correlation with *pyrR* (> 0.9).

**Figure 8.**
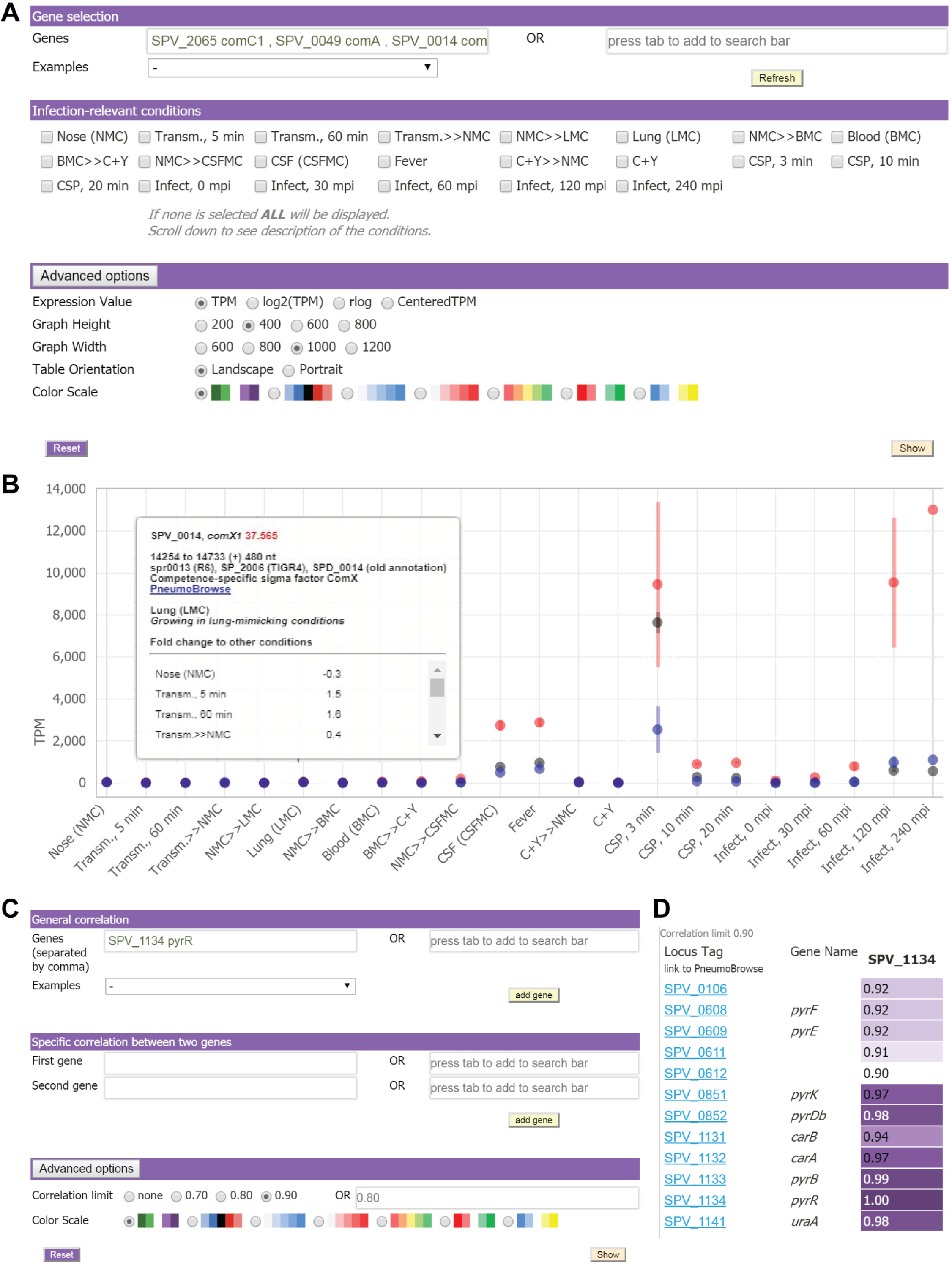
An intuitive, interactive database for accessing expression and correlation data. **A**. Users can specify their gene(s) of interest in the field ‘Genes’. Other settings, including normalization method, color scales and graph dimensions, can be adjusted under “Advanced options”. Multiple genes of interest are queried separated by commas. The immediate genomic environment of the gene(s) of interest can be explored in PneumoBrowse by clicking the locus tag in the result table. **B**. Target expression values are plotted against infection-relevant conditions, and the values can be downloaded for further analysis. The example shown consists of three competence genes. Hovering on a point reveals more information. **C**. The co-expression matrix can be mined by a simple inquiry of a gene of interest (General correlation) while specific correlation provides the correlation value between two genes of interest. Additionally, users can specify a desired threshold for co-expression values under “Advanced options”. **D**. Correlation values to *pyrR*, noting that self-correlation is 1.

## DISCUSSION

Extensive mineable transcriptome databases exist only for a few model bacteria, such as *B. subtilis* (32, 84), *S. aureus* (33, 85), *Escherichia coli* (86, 87) and *Salmonella enterica* serovar Typhimurium (88), and have been proven to be invaluable for the research community. Here, we set out to map the transcriptomic landscape of the important opportunistic human pathogen *S. pneumoniae*. In this study, we coupled exposure to wide-ranging and dynamic infection-relevant conditions (**Table 2** and **Figure 1A**) with high-throughput RNA-seq and generated a compendium of the pneumococcal transcriptome. This use of infection-relevant conditions is similar to what has been successfully applied to other bacteria, including *B. subtilis* (32), *S. enterica* (35), *S. aureus* (33) and *Helicobacter pylori* (34) to incite genome-wide transcriptional responses under wide-ranging physicochemical conditions. Our work highlights key facts about the survival techniques utilized by *S. pneumoniae*, such as the substantial transcriptional regulation of sugar transporters (**Figure 5A**, **Supplementary Tables S3** and **S4** on ‘Dynamic Genes’), mainly in response to the presence of alternative sugars or in the absence of glucose and mediated by the transcription factor CcpA. These observations indicate the necessity of acquiring a carbon source for pneumococcal in-host survival as shown in several *in vivo* experiments (64, 89).

Exposure to conditions relevant to the natural lifestyle of various bacteria has been reported to incite genome-wide transcriptional responses (32, 34, 90, 91). Here, we show that under a set of varied infection-relevant conditions, there was a subset of genes that was constantly highly expressed while there was no gene that was always lowly expressed – highlighting the saturated and dynamic nature of the pneumococcal transcriptome (**Figures. 2B, 3C** and **5A**). Previously, we reported that all pneumococcal genes were expressed during early infection (26), and this was again confirmed in this study because none of the genes were consistently silent.

The pneumococcus occupies a rich and diverse niche of the respiratory tract (13). While we tried to estimate the relevant conditions for the pneumococcus during its pathogenic lifestyle, other important physicochemical parameters which we did not include in the infection models, such as the concentration of metal ions, play important roles in survival (92) and virulence (93). Moreover, the pneumococcus shares a busy ecosystem in the respiratory tract with other bacteria, fungi and viruses (13). Activities of other residents may be detrimental to the pneumococcal survival, as in the case of *Haemophilus influenzae* recruiting host cells to remove *S. pneumoniae* (94). On the other hand, pneumococcal interactions with influenza viruses yield bountiful nutrients to support pneumococcal expansion (95). Dual transcriptomics studies involving the interaction with other relevant species will offer interesting insights into pneumococcal gene expression and will greatly enhance our understanding of pneumococcal biology and pathogenesis (26, 96).

Additionally, we have proposed a simple and straightforward manner for converting the dense and substantial sequencing data into a type of gene network that we call the co-expression matrix (**Figure 6**). The matrix was assembled by arranging correlation values between two genes by their respective genomic locations, and its potential was demonstrated by the elucidation of a new member of the ComE regulon, called *briC* (SPV_0391, **Figure 7**), indicating that it can be a valuable tool for developing new hypotheses regarding cellular pathways or gene functions. Nevertheless, downstream experiments should be performed to verify these hypotheses (97). Lastly, we provide the comprehensive and rich dataset to the research community by building a user-friendly online database, PneumoExpress (https://veeninglab.com/pneumoexpress), where users can easily extract expression values and fold changes of a gene of interest, as well as quantitative information on how its expression profile correlates with that of other genomic features (**Figure 8**). By a simple click in the database, users can explore the immediate genomic environment of genes of interest in PneumoBrowse (28). In addition, the resources assist efforts in comparative genomics and transcriptomics for other bacteria. Finally, we invite other researchers to harness these resources and generate their own hypotheses to gain new insights into pneumococcal biology and, ultimately, to identify novel treatment and prevention strategies against pneumococcal disease.

## DATA AVAILABILITY

The source code for the online compendium is available in Zenodo, https://doi.org/10.5281/zenodo.1293939. Licensed under Creative Commons Attribution-Non Commercial. The transcriptomic datasets are available in the GEO repository: accession number GSE108031. PneumoExpress is hosted on two separate servers: http://pneumoexpress.molgenrug.nl and https://veeninglab.com/pneumoexpress-app.

## ACKNOWLEDGEMENTS

We are grateful to V. Benes and B. Haase (GeneCore, EMBL, Heidelberg) for their continuing support in sequencing; C.J. Albers, B. Jayawardhana, E.C. de Wit and M.H. Silvis for many fruitful discussions; A. de Jong for bioinformatics support; and A. Lun (Cancer Research UK, Cambridge) for insightful recommendations concerning rRNA analysis. We would like to thank the Center for Information Technology of the University of Groningen for their support and for providing access to the Peregrine high-performance computing cluster. We appreciate the following creators: Hyhyhehe, Misha Petrishchev, Alberto Gongora, Hea Poh Lin, and Icon 54 for making their cliparts freely available at thenounproject.com.

## FUNDING

Work in the Veening lab is supported by the Swiss National Science Foundation (project grant 31003A_172861), a VIDI fellowship (864.11.012) of the Netherlands Organization for Scientific Research (NWO-ALW), a JPIAMR grant (50-52900-98-202) from the Netherlands Organisation for Health Research and Development (ZonMW) and ERC starting grant 337399-PneumoCell.

## CONFLICT OF INTEREST

The authors declare that they have no competing interests.

