## Supplementary Materials for "High-resolution analysis of the pneumococcal transcriptome under a wide range of infection-relevant conditions" (from another version of this preprint: version 2, 2018-03-22)

**SUPPLEMENTARY INFORMATION**

**Supplementary Figures**

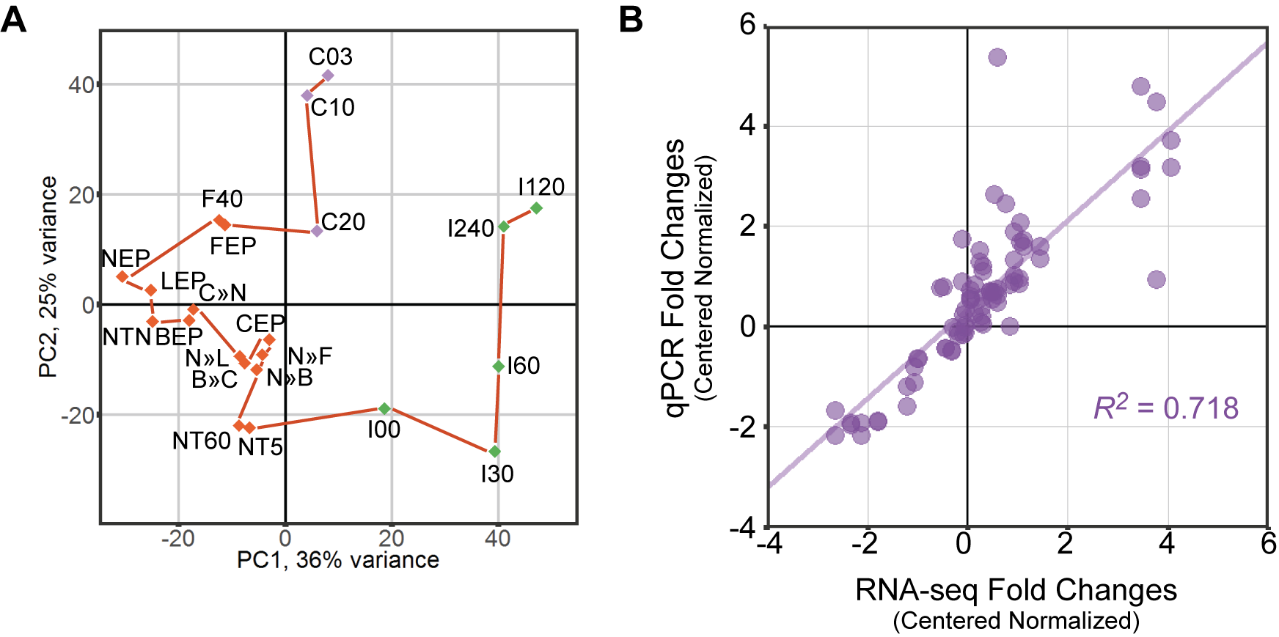

**Supplementary Figure S1**. **A**. The shortest tour between infection-relevant conditions. The tour was calculated by the TSP algorithm on a two-dimensional plane based on an Euclidean distance. **B**.  Fold changes of qPCR validates fold changes obtained from RNA-seq. The values were normalized and centered according to the technology used.

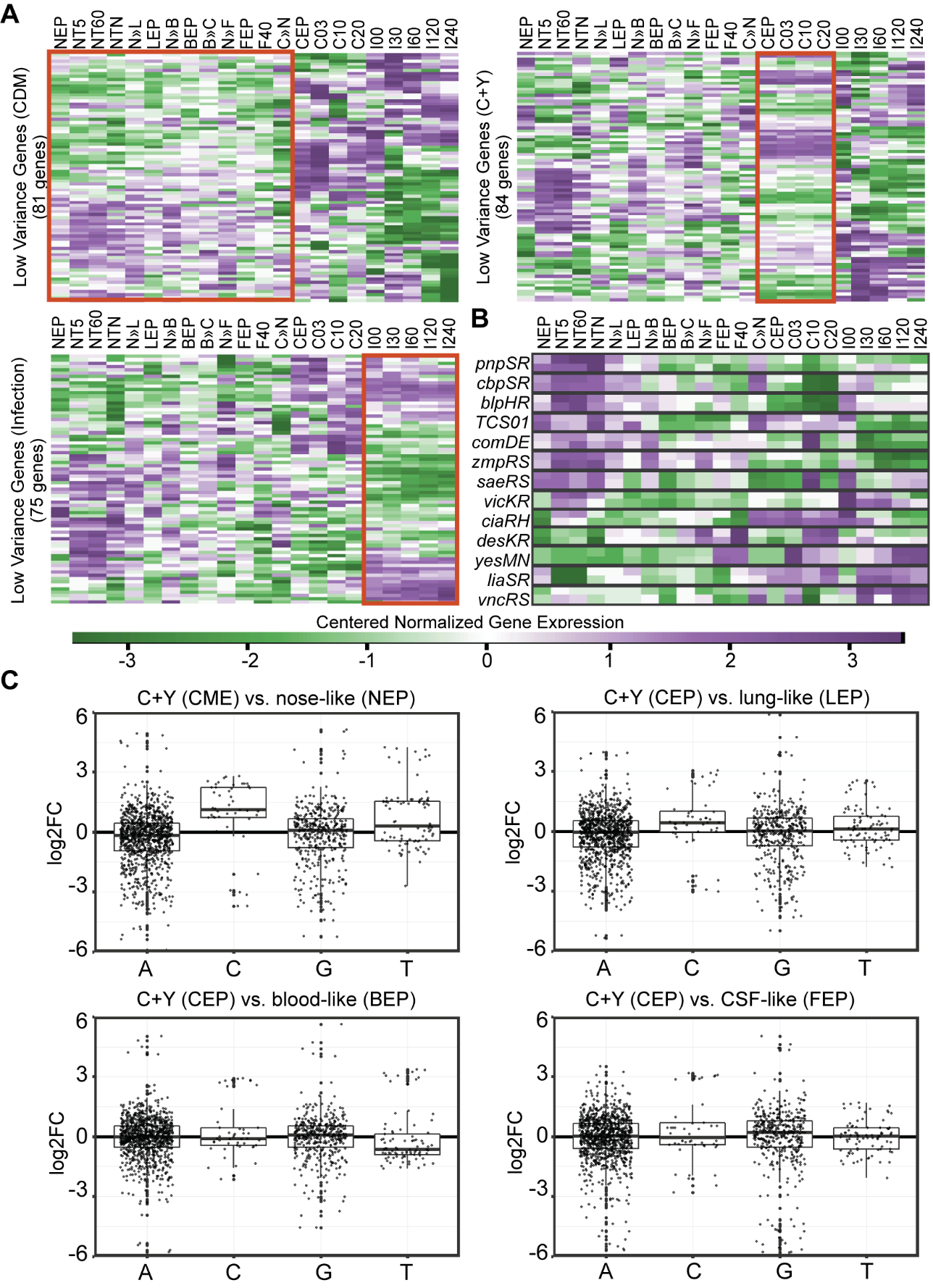

**Supplementary Figure S2**. **A**. Normalized gene expression was centered across infection-relevant conditions and shown as heatmap. Subsets of low variance genes in each basal mediums: CDM, C+Y and infection medium. Orange boxes indicates specific conditions based on each medium. **B**. Centered normalized expression of two component systems (TCSs) across infection-relevant conditions. Purple indicates higher expression while green low expression. **C**. Fold changes of operons in C+Y, which support the highest growth rate vs. infection-like sites separated by the starting nucleotide of each operon. Nose-like medium support the lowest growth rate, followed by lung-like, blood-like and CSF-like mediums.

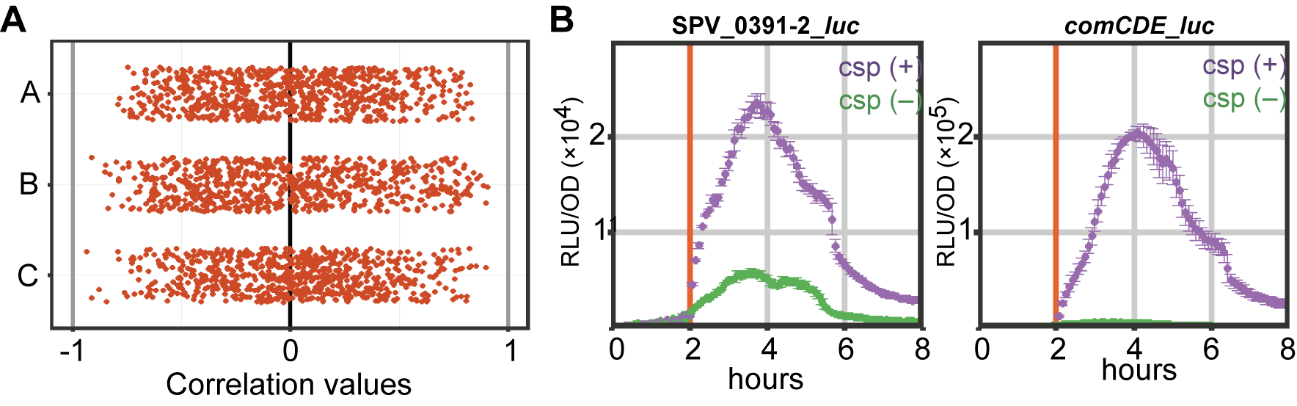

**Supplementary Figure S3**. **A**. Correlation values of three cluster containing 40 features randomly selected from pneumococcal annotated features. **B**. Luminescence signal from firefly luciferase transcriptionally fused to SPV_0391 and *comCDE* due to addition of exogenous csp-1, competence stimulating peptide-1, 2 hours after incubation (orange line).

**Supplementary Materials and Methods**

**Culturing of *Streptococcus pneumoniae* D39 and transformation**

*S. pneumoniae* strains were routinely cultured from frozen stock in C+Y (pH 6.8), a liquid casein-based medium, supplemented with yeast extract (Sigma-Aldrich, Boom, NL) at 37°C. Prior to culturing for RNA isolation, *S. pneumoniae* was pre-cultured in the corresponding CDMs.

Construction of strains was performed as previously described (1). For transformation, cells were grown until OD_600nm_ ~0.1. Subsequently, synthetic CSP-1 (100 ng·ml^-1^**,** competence-stimulating peptide-1) was added into the suspension and incubation was continued for another 12 min at 37°C. Transforming DNA was added to the now-competent *S. pneumoniae*. Cells were then incubated at 30°C for 20 min to allow for DNA internalization. Afterwards, fresh C+Y was added and the suspension was incubated further at 37°C for 90 minutes. Selection of transformants was done by plating in Columbia agar (Oxoid, UK) supplemented with 2% sheep blood (Johnny Rottier, NL) and 0.25 mg·ml^-1^ kanamycin for *luc* constructs, 0.05 μg·ml^-1^ erythromycin for deletion constructs. The following day, colonies were streaked on antibiotic-supplemented blood agar and incubated overnight in 5% (v/v) CO_2_. Constructs were checked by colony PCR and Sanger sequencing.

**Preparation of stock solutions for modified Sicard’s Chemically Defined Media**

The Sicard’s media were prepared from pre-prepared stock solutions of: basal solution, amino acids, micronutrients, nucleobases, vitamins, pyruvate, choline, bovine serum albumin (BSA), glucose and glucosamine. 1 L of nose- and lung-like basal solution contained 18 ml K_2_HPO_4_ 1 N, 28 ml KH_2_PO_4_ 1 N, 750 mg ammonium citrate, 1.25 g sodium acetate and 500 mg cysteine-HCl. 1 L of blood-like basal solution contained 134 ml K_2_HPO_4_ 1 N, 33 ml KH_2_PO_4_ 1 N, 1 g ammonium citrate, 1.67 g sodium acetate and 667 mg cysteine-HCl. 1 L of CSF-like basal solution contained 114 ml K_2_HPO_4_ 1 N, 12 ml KH_2_PO_4_ 1 N, 750 mg ammonium citrate, 1.25 g sodium acetate and 500 mg cysteine-HCl.

1 L amino acid solution contained 3 g L-alanine, 1.55 g L-arginine, 4.4 g L-asparagine, 5.25 L L-aspartate, 6.25 L-glutamate, 4.9 g L-glutamine, 2.2 g glycine, 1.9 g L-histidine, 2.65 g L-isoleucine, 5.7 g L-leucine, 5.5 g L-lysine, 1.55 g L-methionine, 3.45 g L-phenylalanine, 8.45 g proline, 4.25 g L-serine, 2.8 g L-threonine, 0.65 g L-tryptophan and 4.05 g L-valine - dissolved and pH-adjusted to 6.5. The solution was then filtered (0.22 µm), aliquoted and kept at -20°C, away from light.

1 L micronutrient solution contained 20 g MgCl_2_, 3.8 g CaCl_2_ and 500 mg ZnSO_4_. The solution was then filtered (0.22 µm) and kept at 4°C. 1 L nucleobases solution contained 1 g adenine, 1 g uracil, 1 g xanthine and 1 g guanine, dissolved in 0.1 N NaOH, filtered (0.22 µm) and kept at 4°C. 10% pyruvate, 2.5 g·l^-1^ choline, 25% BSA, 20% glucose and 20% glucosamine solutions were prepared, filtered (0.22 µm) and kept at 4°C.

1 L vitamin solution contained 500 mg sodium aminobenzoate, 250 mg biotin, 100 mg folic acid, 100 mg nicotinic acid, 100 mg calcium pantothenate, 250 mg pyridoxamine-HCl, 200 mg pyridoxine-HCl, 100 mg riboflavin, 100 mg thiamine-HCl, 150 mg lipoic acid and 100 mg vitamin B12. Riboflavin was dissolved in around 750 ml mQ and heated up to 70°C. Other vitamins were added when the solution was around 30°C. Biotin and folic acid were dissolved in 2 N NaOH, added into the solution and pH-adjusted to 6.5, filtered (0.22 µm) and kept at 4°C. Unless otherwise stated, all of the chemicals were supplied by Sigma-Aldrich (Boom, NL).

100 ml nose-like medium was made by sterilely mixing 80 ml nose/lung basal solution, 8 ml amino acids, 1 ml micronutrients, 1 ml nucleobases, 1 ml vitamins, 0.1 ml pyruvate, 0.8 ml choline, 0.4 ml 25% BSA and 0.64 ml glucosamine solutions. pH was adjusted to 7.0 and sterile mQ was added up to 100 ml. 100 ml lung-like medium was made by sterilely mixing 80 ml nose/lung basal solution, 8 ml amino acids, 1 ml micronutrients, 1 ml nucleobases, 1 ml vitamins, 0.1 ml pyruvate, 0.8 ml choline, 1.2 ml 25% BSA and 0.32 ml glucosamine solutions. pH was adjusted to 7.0 and sterile mQ was added up to 100 ml. 100 ml blood-like medium was made by sterilely mixing 60 ml blood basal solution, 8 ml amino acids, 1 ml micronutrients, 1 ml nucleobases, 1 ml vitamins, 0.1 ml pyruvate, 0.8 ml choline, 26.8 ml 25% BSA and 0.45 ml glucose solutions. pH was adjusted to 7.4 and sterile mQ was added up to 100 ml. Lastly, 100 ml CSF-like medium was made by sterilely mixing 80 ml CSF basal solution, 8 ml amino acids, 1 ml micronutrients, 1 ml nucleobases, 1 ml vitamins, 0.1 ml pyruvate, 0.8 ml choline, 0.18 ml 25% BSA and 0.225 ml glucose solutions. pH was adjusted to 7.8 and sterile mQ was added up to 100 ml.

**Preparation of stock solutions for C+Y**

The C+Y medium is prepared from pre-prepared stock solutions of: Pre-C, Adams solutions, yeast extract, HCl, sodium pyruvate, bovine serum albumin (BSA), uridine and adenosine, glucose, MnCl_2_, glutamine and sucrose. 1L of Pre-C solution contains 11.25 mg cysteine-HCl, 2 g sodium acetate, 5 g casein hydrolysate, 6 mg L-tryptophan and 8.5 g K_2_HPO_4_ - dissolved and sterilized by autoclaving (121°C, 15 min), kept at room temperature, away from light.

1L Adams I solution contains 150 mg biotin, 150 mg nicotinic acid, 175 mg pyridoxine-HCl, 600 mg calcium pantothenate, 160 mg thiamine-HCl and 70 mg riboflavin - dissolved and pH-adjusted to 7.0 by 1N solution of NaOH. The solution was then sterilized in boiling water (15 min). 1L Adams II solution contains 5 g FeSO_4_.7H_2_O, 5 g CuSO_4_.5H_2_O, 5 g ZnSO_4_.7H_2_O and 2 g MnCl_2_.4H_2_O - dissolved in 900 ml mQ and 100 ml fuming HCl (37%, 11.4 N). The solution was sterilized in boiling water (15 min). 1L Adams III solution contains 128 ml Adams I, 3.2 ml Adams II, 1.4 g L-asparagine anhydrous, 160 mg choline, 400 mg CaCl_2_ anhydrous and 16 g MgCl_2_.6H_2_O - dissolved and pH-adjusted to 7.6 using NaOH 10N. The solution was sterilized by filtration (0.22 µm). All Adams solutions were kept at 4°C, away from light.

Yeast extract solution (10%) was prepared by suspending yeast extract in mQ and autoclaving (121°C, 15 min). The solution was then filtered (0.22 µm), aliquoted and kept at -20°C, away from light. 1 N HCl is prepared from fuming HCl and further filtered (0.22 µm) and kept at 4°C. 2% sodium pyruvate solution, 8% BSA solution, 0.4 mM MnCl_2_ solution, 20% glucose and 1.5 M sucrose solutions were prepared, filtered (0.22 µm) and kept at 4°C. Uridine-adenosine solution was prepared by dissolving 2 mg·ml^-1^ uridine and 2 mg·ml^-1^ adenosine, filtered (0.22 µm) and kept at 4°C. 3% glutamine was prepared, filtered (0.22 µm) and kept at -20°C. Unless otherwise stated, all of chemicals were supplied by Sigma-Aldrich (Boom, NL).

100 ml acidified C+Y medium was made by mixing 80 ml of Pre-C, 2 ml of Adams III, 2 ml of yeast extract, 1.6 ml HCl 1N, 1.2 ml sodium pyruvate, 0.8 ml 8% BSA solution, 0.8 ml glucose, 0.8 ml uridine-adenosine, 80 µl MnCl_2_, 58.4 µl glutamine and 50.4 µl sucrose. C+Y was made fresh and stored at 4°C.

**Infection-like pneumococcal cultures**

In order to simulate infection in a laboratory, five types of chemically defined media (CDMs) were prepared. Acidified C+Y medium was prepared fresh without modification. Four media were modified from Sicard's defined medium (2) by variation of the type and quantity of carbon source, level of serum albumin, starting pH, exposure to CO_2_ and growth temperature (**Fig. 1B**). For complete content of the media, *see* **Supplementary Table 1**. Stock cultures (-80°C) of *S. pneumoniae* D39 were inoculated into selected CDMs, grown up to OD_595nm_ ~0.4 at appropriate temperature (30°C for nose-like conditions and 37°C for lung-, blood-, CSF-like and C+Y media), and then re-inoculated 1:100 into a fresh batch of the same CDM. Cultures were harvested for RNA isolation during exponential phase for the five growth conditions (NEP, LEP, BEP, FEP and CEP).

In transfer experiments, cells were harvested from growth culture by spinning down at 4°C (maximum speed, 3 min), removing supernatant and resuspending in target medium for 5 minutes, after which the cells were immediately harvested. In the meningeal fever experiment, a culture in CSF-like medium was transferred to 40°C for 5 minutes and harvested. In competence experiments, 100 ng·ml^-1^ CSP-1 was added into C+Y culture and cells were harvested after 3, 10 and 20 minutes.

In transmission experiments, cells growing in nose-like conditions were spun down (4°C, 3 min) and supernatant was removed. The cell pellet was resuspended with leftover supernatant and pipetted out onto a sterile plastic surface under laminar air flow for 5 (NT5) or 60 minutes (NT60). In order to harvest the cells, saturated ammonium sulfate solution is added directly onto dried bacterial culture and pipetted into a new tube. In NTN, the dessicated cells were resuspended into nose-like medium (30°C, 5 min) before harvesting. Co-incubation cultures were described previously [(17)](https://www.colwiz.com/cite-in-google-docs/cid=f20fb61e2a9750e).

**Total RNA isolation**

Isolation of RNA was performed as previously described with some modifications [(17)](https://www.colwiz.com/cite-in-google-docs/cid=f20fb61e2a9750e). In order to stop protein-dependent RNA production and degradation, we treated the pneumococcal culture with concentrated solution of ammonium sulfate (3). Each milliliter of the ammonium sulfate solution (pH 5.2) contained 0.7 g (NH_4_)_2_SO_4_. The saturated solution also contained 20 mM EDTA and 25 mM sodium citrate. Three parts of saturated solution of ammonium sulfate was added directly to one part of culture, immediately followed by vigorous mixing. The suspension was diluted twice with DEPC-treated mQ before centrifugation at full speed (20 min, 4 °C, 10,000 × g). Supernatant was removed and the pellet was snap-frozen with liquid nitrogen.

In a 1.5 ml screw cap tube, a PCR tube full of sterile, RNase-free glass beads (100 μm, BioSpec, US) were added together with 50 μl 10% SDS and 500 μl phenol-chloroform. The frozen pellet was resuspended in TE solution (10 mM Tris-HCl, 1 mM Na2DTA, pH 8.0, DEPC treated mQ). The cell suspension was added into the screw cap tube and bead-beaten three times, 45 s each. Tubes were immediately placed on ice and centrifuged at full speed at 4°C to separate organic and aqueous phases. The aqueous phase was pipetted out and back-extraction was performed on the organic phase to optimize RNA yield. Phenol was further depleted from the aqueous phase by a round of chloroform extraction. After vigorous vortexing, the mixture was again centrifuged at full speed, 4°C. The aqueous phase was pipetted into a new eppendorf tube and nucleic acids were alcohol-precipitated: 50 µl NaOAc 3M and 1 ml of cold isopropanol were added and mixed thoroughly. The mixture was incubated for at least 30 minutes at -20°C before pelleting by centrifugation (full speed, 4°C). Supernatant was removed gently and the nucleic acid pellet was resuspended in ice-cold 75% ethanol before re-pelleting was performed (full speed, 4°C). Ethanol washing was performed once more. The pellet was air-dried before DNase treatment.

DNase treatment was performed according to the manufacturer’s protocols (RNase-free DNase I recombinant, Roche, US) for 1 hour, room temperature. To remove DNase and gDNA-derived nucleotides, phenol-chloroform extraction, chloroform extraction, isopropanol precipitation and ethanol washing were performed as previously mentioned. Total RNA was resuspended in 30 µl TE buffer. The quantity and quality of total RNA was estimated by Nanodrop and a 1% bleach gel was employed to interrogate the presence of genomic DNA and rRNA bands (23S, 2.9 kbp and 16S, 1.5 kbp).

**Library preparation, sequencing, data pipeline, and online database**

Without depleting ribosomal RNAs, stranded cDNA library preparation was performed for the 34 samples with the TruSeq® Stranded Total RNA Sample Preparation Kit (Illumina, US), according to the manufacturer’s protocol. cDNA sequencing of the 34 samples was performed in six lanes of Illumina NextSeq 500 with a HighOutput Flowcell in 75 nucleotide single-end mode. Samples were de-multiplexed and analyzed further. The raw FASTQ data are accessible at <http://www.ncbi.nlm.nih.gov/geo/> with accession number GSE108031.

The quality of raw reads was checked by FastQC v0.11.5 (Babraham Bioinformatics, UK, (4). To improve the quality of reads and remove adapter sequences, we trimmed the reads using Trimmomatic 0.36 (5) in single-end mode using these criteria: (i) adapter sequences removed based on TruSeq3-SE library; (ii) low-quality leading nucleotides removed; (iii) low-quality trailing nucleotides removed; (iv) truncated reads when the average quality score in a five-nucleotide sliding window dropped below 20; and (v) removed reads shorter than 50 nucleotides. The trimmed reads were again checked by FastQC. As reference genome, we used the in-house generated genome with its corresponding annotation (BioRxiv: <https://doi.org/10.1101/283663>), excluding 264 features that are contained by annotated pseudogenes. To ensure an undisrupted mapping of SPV_0001, a pneumococcal gene located at the very beginning of the pneumococcal genome sequence file, we copied the first 600 bp of the pneumococcal genome to the end of the genome and annotated this extra copy as SPV*_*0001a. Read counts of this extra feature were combined with those of SPV_0001 for downstream analysis. Alignment of trimmed reads to the chimeric genome was performed by STAR (6), with the following options: (i) alignIntronMax 1 and (ii) sjdbOverhang 49. The aligned reads were then counted (7) according to the in-house annotation file in (i) multimapping mode (-M), in which fractional count was reported (--fraction), (ii) allowing for overlapping reads across features to accommodate operons (iii) in a stranded option.

Subsequently, we analyzed the libraries in R-studio (R v3.4.2). We performed differential gene expression analysis on rounded raw count by DESeq2 (8). For visualization of gene expression, the libraries were regularized logarithm-normalized by DESeq2. Aside from DESeq2-transformed counts, the rounded raw counts were transformed into TPM (transcripts per million) and log-transformed TPM (9). Centering of gene expression for visualization was performed by calculating Z-score of every value within the complete set of expression values for a specific gene across all conditions. The three transformed counts are used to visualize gene expression level in our online database (www.veeninglab.com/pneumoexpress). The data are stored in a MySQL database containing *S. pneumoniae* gene expression values. Gene expression graphs during early infection are generated by D3 (Data Driven Documents, [https://d3js.org](https://d3js.org/)).

**Generation of pneumococcal *luc*-fusion and deletion strains**

In order to monitor the dynamics of gene expression, we tagged target genes with a downstream gene encoding firefly luciferase (*luc*). This *luc* insertion was performed based on enzymatic restriction-ligation of three fragments: (i) ~1000 bps upstream of insertion including the target gene, (ii) *luc*-kanamycin-resistance cassette and (iii) ~1000 bps downstream of insertion. The *luc* gene retained its ribosome-binding site (RBS) while the kanamycin resistance marker is under control of its own promoter. For deletion strains, the same strategy was employed with the only difference being fragment 2 was an erythromycin-resistance cassette under its own promoter. Assembled fragments were transformed into the wild-type strain. All oligonucleotides involved in fragment construction are listed in **Supplementary Table 10** while pneumococcal strains are listed in **Supplementary Table 11**.

**Luminescence assays**

Assays were performed as described previously (10) with minor changes. *S*. *pneumoniae* was pre-cultured to OD_600_ 0.2 and diluted 100-fold in C+Y medium containing 250 μg·ml^-1^ D-luciferin prior to the assay. Production of firefly luciferase (encoded by *luc*) causes emission of light when the substrate, D-luciferin is available in the medium. Luminescence assays were performed in 96-well clear bottom white plates (Costar®, US) at 37°C. OD_595_ and luminescence (as relative luminescence units, RLU) were measured every 5 minutes using a Tecan Infinite 200 PRO instrument.

**qRT-PCR as confirmation of RNA-sequencing**

Total RNA isolation, reverse transcription and qPCR were performed as previously described (1). cDNA was synthesized from total RNA by SuperScript® III Reverse Transcriptase (Life Technologies, NL). The nucleic acid mix contained total RNA sample, random nonamers, dNTP mix (Life Technologies, NL) and RiboLock RNase Inhibitor (Life Technologies, NL). The nucleic acid mix was then incubated at 65°C for 5 minutes and put directly on ice. First-strand buffer, DTT, RiboLock and Reverse Transcriptase were added into the mix and incubated at 25°C for 5 min, 50°C for 60 min and 70°C for 15 min.

Amplification efficiency for each primer pair was calculated based on its ability to double the amount of product per cycle. The qPCR mix contained forward and reverse primers, SYBR Green Real-Time PCR Master Mix (Life Technologies, NL) and cDNA – the reactions were performed in triplicate and at two different cDNA concentrations. The PCR temperature scheme was as follows: 95°C, 3 min, followed by 40 cycles of 95°C, 10 s; 50°C, 30 s and 72°C, 30 s on an iQ™5 Optical System (BioRad, NL). Data were analyzed according to the ∆∆Ct method (11), against housekeeping gene, *gyrA*. Gene expression values were then normalized prior to comparison with data generated by sequencing. Fold change was calculated against one sample: growth in nose-like (NEP).
